# Cell-cycle dependent inhibition of BRCA1 signaling by the lysine methyltransferase SET8

**DOI:** 10.1101/2023.11.22.567520

**Authors:** Yannick Perez, Fatima Alhourani, Julie Patouillard, Cyril Ribeyre, Marion Larroque, Véronique Baldin, David Lleres, Charlotte Grimaud, Eric Julien

## Abstract

The cell-cycle regulated methyltransferase SET8 is the sole enzyme responsible for the mono-methylation of histone H4 at lysine 20 (H4K20) that is the substrate for di- and tri-methylation mainly by SUV4-20Hs enzymes. Both SET8 and SUV4-20Hs have been implicated in regulating DNA repair pathway choice through the inverse affinities of BRCA1-BARD1 and 53BP1 complexes for disparate methylation states of H4K20. However, the precise and respective functions of each H4K20 methyltransferases in DNA repair pathways remained to be clarified. Here, we show that SET8 acts as a potent chromatin inhibitor of homologous recombination and that its timely degradation during DNA replication is essential for the spontaneous nuclear focal accumulation of BRCA1 and RAD51 complexes during S-phase. Strikingly, the anti-recombinogenic function of SET8 is independent of SUV4-20H activity but requires the subsequent recruitment of the ubiquitin ligase RNF168. Moreover, we show that SET8-induced BRCA1 inhibition is not necessarily related to the loss of BARD1 binding to unmethylated histone H4K20. Instead, it is largely caused by the accumulation of 53BP1 in a manner depending on the concerted activities of SET8 and RNF168 on chromatin. Conversely, the lack of SET8 and H4K20 mono-methylation on newly assembly chromatin after DNA replication led to the untimely accumulation of BRCA1 on chromatin at the subsequent G1 phase. Altogether, these results establish the *de novo* activity of SET8 on chromatin as a primordial epigenetic lock of BRCA1-mediated HR pathway during the cell cycle.

## INTRODUCTION

The genome of eukaryotic cells is constantly challenged by variety of DNA insults, which can occur in response to exogenous agents or accidentally during DNA-dependent processes. Double-strand breaks (DSBs) represent the most detrimental lesions, because failure to eliminate them can lead to cell death or a wide variety of genetic alterations (Xu and Xu 2020). To deal with this threat, cells have evolved an elaborate DNA repair network where homologous recombination (HR) and non-homologous end joining (NHEJ) are the two major pathways used for repairing DSBs [2,3]. In contrast to NHEJ, HR usually depends on the presence of a sister chromatid produced by DNA replication, as it provides an undamaged homologous template for repair of both broken strands. The use of HR is therefore closely linked to S and G2 phases of the cell cycle [3]. Furthermore, most of HR factors are naturally found enriched in the vicinity of replication forks, suggesting that they play important role in the stability of genome during S-phase and making HR as the favored repair pathway to deal with replication-associated DNA lesions [4]. However, while the cascade of events that lead to the activation of homologous recombination is extensively studied in the context of exogenous DNA lesions, less is known about the mechanisms that regulate HR during unperturbed S-phase progression. Notably, how HR signaling pathway is specifically restrained to both S and G2 phases of the cell cycle is still poorly understood. Yet, these mechanisms are likely essential, since inaccurate activation or misuse in HR pathways during the other phases of the cell cycle can cause deleterious outcomes associated with cancer [5,6].

A critical determinant for the regulation of HR signaling pathway is the balanced activities between the pro-NHEJ protein 53BP1 and the HR-promoting heterodimer formed by BRCA1 and BARD1 during the cell cycle [2,3]. Upon DNA damage, 53BP1 accumulates at DSB sites by recognizing notably nucleosomes marked by the di-methylation of histone H4 at lysine 20 (H4K20me2) and to lesser extent ubiquitylation of histone H2A at lysine 15 [7]. This 53BP1 accumulation on damaged chromatin limits HR by antagonizing the DNA end resection activity of BRCA1-BARD1 and thus the subsequent loading of RAD51 that catalyzes the repair reaction by recombination [8]. As cells progress through S phase, the ability of 53BP1 to accumulate around DNA breaks decline. This has been notably attributed to the replication-coupled dilution of H4K20me2 mark on post-replicated chromatin [8,9]. Yet, 53BP1 can also interact with histone H4K20 mono-methylation (H4K20me1) and tri-methylation (H4K20me3) [10,11], thereby questioning this model. Conversely, the lack of methylation of H4K20 (H4K20me0) is a mark of post-replicative chromatin specifically recognized by the ankyrin domain of BARD1, which contributes to BRCA1 recruitment and the eviction of 53BP1 when sister chromatid is available for HR repair [12,13]. Thus, the inverse affinities of 53BP1 and BARD1 for distinct methylation sates of histone H4K20 have underscored a potential key role of this lysine methylation in the regulation of DNA repair pathways choice during the cell cycle. However, this hypothesis has been explored so far only in response to exogenous DNA damaging agents and never been validated in a more physiological and unchallenged context. Furthermore, the roles of H4K20 enzymes responsible and the biological significance of different H4K20me states in the cell-cycle regulatory interplay between 53BP1 and BRCA1 still remain to be clarified.

SET8 is the sole enzyme responsible for the mono-methylation of histone H4 at lysine 20 (H4K20) that is the substrate for di- and tri-methylation by SUV4-20H enzymes. In this work, we demonstrate that SET8 functions as a potent inhibitor of homologous recombination at the chromatin level, and its timely degradation during DNA replication is crucial for the spontaneous accumulation of BRCA1 and RAD51 complexes in the vicinity of replication forks. Independent of SUV4-20H activity, the anti-recombinogenic role of SET8 involves the recruitment of the ubiquitin ligase RNF168, resulting in a post-replicated chromatin state that hinders BRCA1-dependent homologous recombination outside of S and G2 phases. This is not necessarily related to the loss of BARD1 binding to H4K20me0, but rather to the accumulation of 53BP1 on post-replicated chromatin in a manner depending on the concerted activities of SET8 and RNF168.

## RESULTS

### PCNA-induced SET8 degradation triggers H4K20me0 enrichment on post-replicated chromatin

Previous studies showed that the newly incorporated histones H4 during S-phase remained unmethylated at lysine 20 (H4K20me0) until the onset of mitosis [14]. The absence of methylation of newly synthetized histone H4 may be related to the higher activity of H4K20 demethylases at the onset of DNA replication, as suggested by previous studies on PHF8 and hHR23 enzymes [15,16]. An alternative but not mutually exclusive possibility raised by other studies [12,17] is that the H4K20me0 enrichment during S-phase is related to the replication-coupled degradation of the lysine methyltransferase SET8 responsible for H4K20 mono-methylation (H4K20me1), a prerequisite for higher H4K20me states (H4K20me2/me3) induced by SUV4-20H enzymes [18–20]. To validate this second possibility, we used an U2OS cell line harboring a tetracycline(Tet)-inducible FLAG-SET8^PIPmut^ enzyme unable to interact with PCNA and thus resistant to the protein destruction mediated by the CRL4^CTD2^ complex during S-phase [21–23]. The advantage of Tet-inducible SET8 cell line is that it allows us to efficiently synchronize and study the immediate impact of FLAG-SET8^PIPmut^ expression on replicated chromatin without triggering the re-replication and DNA damage phenotypes associated with long-term SET8 stabilization, as previously described [18,23]. Thus, U2OS cells were first synchronized at G1/S transition by thymidine block and then the expression of FLAG-SET8^PIPmut^ was induced by tetracycline treatment for at least 2 hours before release into S-phase. Although displaying a slight delay in S-phase entry, tetracycline-treated cells then progressed similarly in S-phase compared to untreated cells as observed by FACS analysis (Figure 1A). As cells progressed into S-phase, immunoblot analysis showed that the levels of endogenous SET8, but not of FLAG-SET8^PIPmut^ mutant, declined rapidly as expected (Figure 1B). Noted of, the apparent stronger downregulation of endogenous SET8 in replicating cells after tetracycline exposure was likely related to the accumulation of FLAG-SET8^PIPmut^ and the slight delay in S-phase progression compared to untreated conditions. Consistent with previous studies [14,23,24], the lack of SET8 in control cells was accompanied by the decrease in the steady state levels of H4K20me1 and the concomitant accumulation of unmethylated H4K20 mark (H4K20me0) (Figure 1B, left panels). In contrast, the expression of the non-degradable FLAG-SET8^PIPmut^ mutant was sufficient to prevent both the reduction in H4K20me1 levels and the appearance of H4K20me0 in tetracycline-treated cells (Figure 1B, right panels). Since H4K20me0 is a hallmark of post-replicated chromatin, FLAG-SET8^PIPmut^ expression is likely sufficient to trigger *de novo* H4K20me1 on newly incorporated histones H4 in the wake of replication forks. To confirm this hypothesis, asynchronous control and tetracycline-treated U2OS cells were pulse-labelled with EdU for 20 minutes and treated with formaldehyde to cross-link protein-DNA complexes. After covalent linkage of biotin-azide to EdU using click chemistry, EdU-labelled DNA and associated proteins were isolated by iPOND (Isolation of Proteins on Nascent DNA) method and then analyzed by immunoblotting [25]. The procedure without click-it reaction served as negative controls. As shown in Figure 1C, immunoblot analysis showed similar levels of histones and PCNA in untreated and tetracycline-treated U2OS samples after click-it reaction, thereby indicating that replisome and chromatin proteins associated with EdU-labelled DNA were efficiently and similarly captured in control and FLAG-SET8^PIPmut^ expressing cells. In these conditions, analysis of H4K20me states by immunoblot revealed a loss of H4K20me0 with a concomitant gain of H4K20me1 upon FLAG-SET8^PIPmut^ binding to nascent chromatin, while the levels of H4K20me2 and H4K20me3, which likely corresponded to recycled histones H4 [14], remained largely unchanged (Figure 1C). Altogether, these results demonstrate that the expression of a non-degradable SET8 enzyme is sufficient to prevent H4K20me0 accumulation by inducing its premature conversion to H4K20me1 on newly incorporated histones H4. Hence, the natural enrichment of H4K20me0 on post-replicated chromatin mainly depends on the replication-coupled degradation of SET8 in U2OS cells.

**Figure 1:**
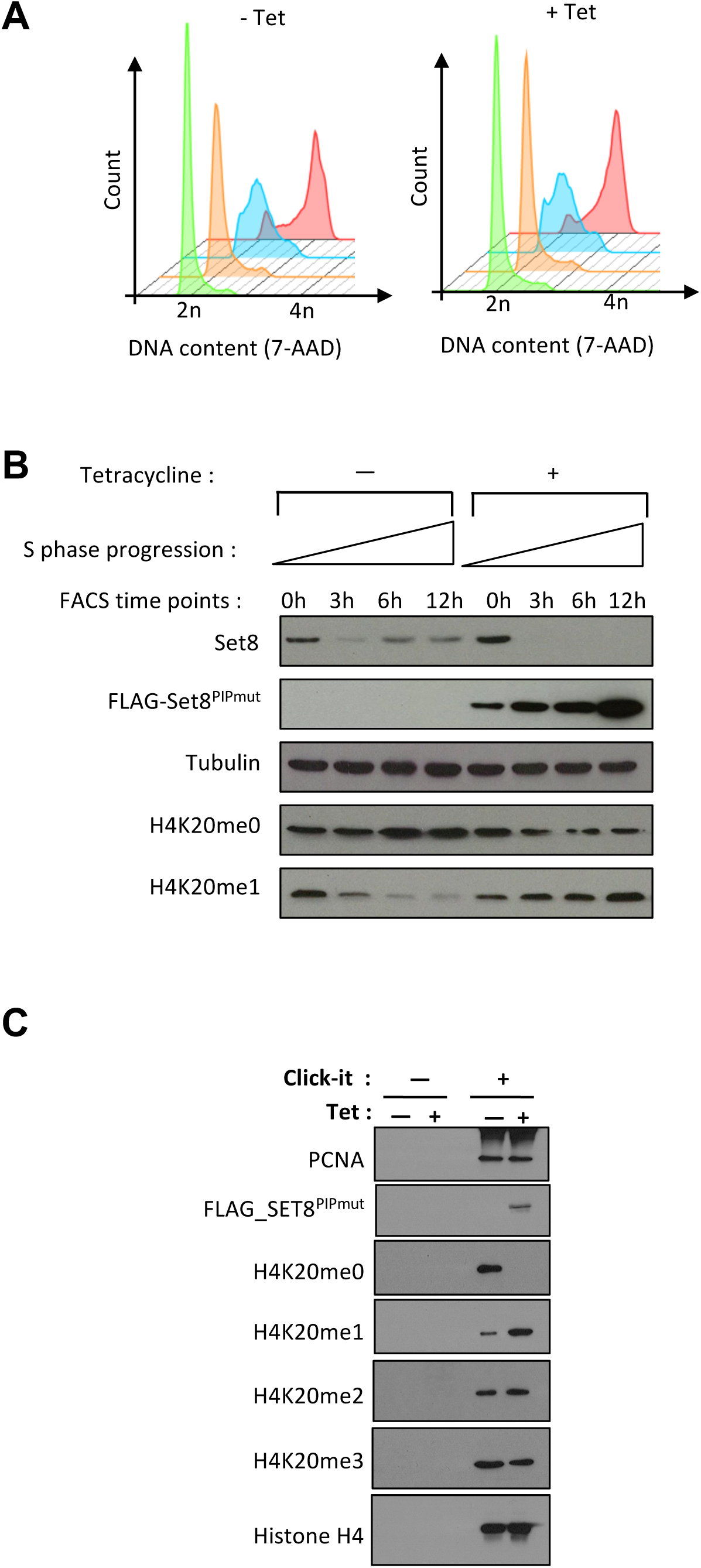
SET8 stabilization induces the switch from H4K20me0 to H4K20me1 on replicated chromatin. **A**. FACS profiles of untreated (−Tet) and tetracycline (+Tet) treated cells synchronized at G1/S transition by double-thymidine block (green profile) and then 3 hours (orange profile), 6 hours (blue profile) and 12 hours (red profile) after released into S-phase. (**B**) Immunoblot analysis of untreated (−Tet) and tetracycline treated (+Tet) cells at each time points analyzed by FACS as shown in Figure 1A. (**C**) Immunoblot analysis of iPOND samples with and without Click-it reaction in untreated and Tet-treated cells. The samples without Click-it reaction serves as negative control of pull-down samples. PCNA and Histone H4 immunoblots serve as loading controls. Similar protein levels (input) before IPOND purification have been verified by immunoblots.

### The replication-coupled functions of BRCA1 are repressed by the presence of active SET8 methyltransferase on chromatin

The recognition of histone H4K20me0 by the BRCA1-associated protein BARD1 allows to recruit BRCA1 and curb 53BP1 anti-resection activity at double-stranded breaks upon ionizing radiation [12]. However, even in absence of exogenous DNA damage, BARD1-BRCA1 complex spontaneously form nuclear foci during S-G2 phases in order to support replication machinery and to ensure genome stability [4,26]. Afterward, the HR functions of BRCA1 and associated proteins must be restrained to avoid genetic aberrations [3]. To explore further the role of SET8 activity and histone H4K20 in the regulation of BRCA1 functions during the cell cycle, we used the Tet-inducible FLAG-SET8^PIPmut^ cellular system and examined by indirect immunofluorescence how the premature conversion of H4K20me0 to H4K20me1 impact on the focal accumulation of BRCA1-BARD1 and localization of 53BP1 in cells progressing normally into S-phase. We also examined the foci formation of the recombinase protein RAD51, as a marker of the activity of BRCA1 during DNA replication [27]. After short treatment or not with tetracycline to induce FLAG-SET8^PIPmut^ expression, cells were pulse-labelled with EdU for 30 minutes before fixation. Fixed cells were then treated with click-it chemistry to reveal EdU incorporation in replicating cells and then co-stained with BARD1, BRCA1, RAD51 or 53BP1 antibodies. The representative images of the results are shown in Figure 2A and their quantifications are shown in Figures 2B and in Supplementary Figure S1A. Consistent with normal S-phase progression (Figure 1A), the pattern of EdU staining was similar between the control and FLAG-SET8^PIPmut^ replicating cells. As expected, BARD1, BRCA1 and RAD51 nuclear foci were detected in control replicating cells (Figure 2A and Figure S1A). We also noticed that these control cells displayed a few and small dispersed nuclear 53BP1 foci (Figure 2A). In contrast, BARD1, BRCA1 and RAD51 nuclear foci were almost abolished in replicating cells upon FLAG-SET8^PIPmut^ expression, whereas the number and size of 53BP1 nuclear foci were strongly increased (Figures 2A and 2B, Figure S1B). Noted of, these nuclear foci alterations were not associated with a specific staining pattern of EdU, suggesting they are not specific to early, mid or late stage of S-phase. To determine whether it depends on the catalytic activity of SET8^PIPmut^, U2OS cells were transduced with pBabe retroviral vectors encoding the active FLAG-SET8^PIPmut^ or its catalytic dead version FLAG-SET8^PIPmut+SETmut^. Three days after retroviral infection, the number of replicating cells displaying RAD51 or 53BP1 nuclear foci was examined by immunofluorescence. As shown in Figures 2C and 2D, the expression of the inactive FLAG-SET8^PIPmut+SETmut^, but not of FLAG-SET8^PIPmut^, failed to reduce RAD51 foci formation and to stimulate 53BP1 focal accumulation during S-phase. Altogether, these results provide the demonstration that the replication-coupled focal accumulation of BRCA1 and its downstream effector RAD51 depends on the cell-cycle regulated timing of SET8 activity on chromatin irrespective to exogenous DNA damage.

**Figure 2:**
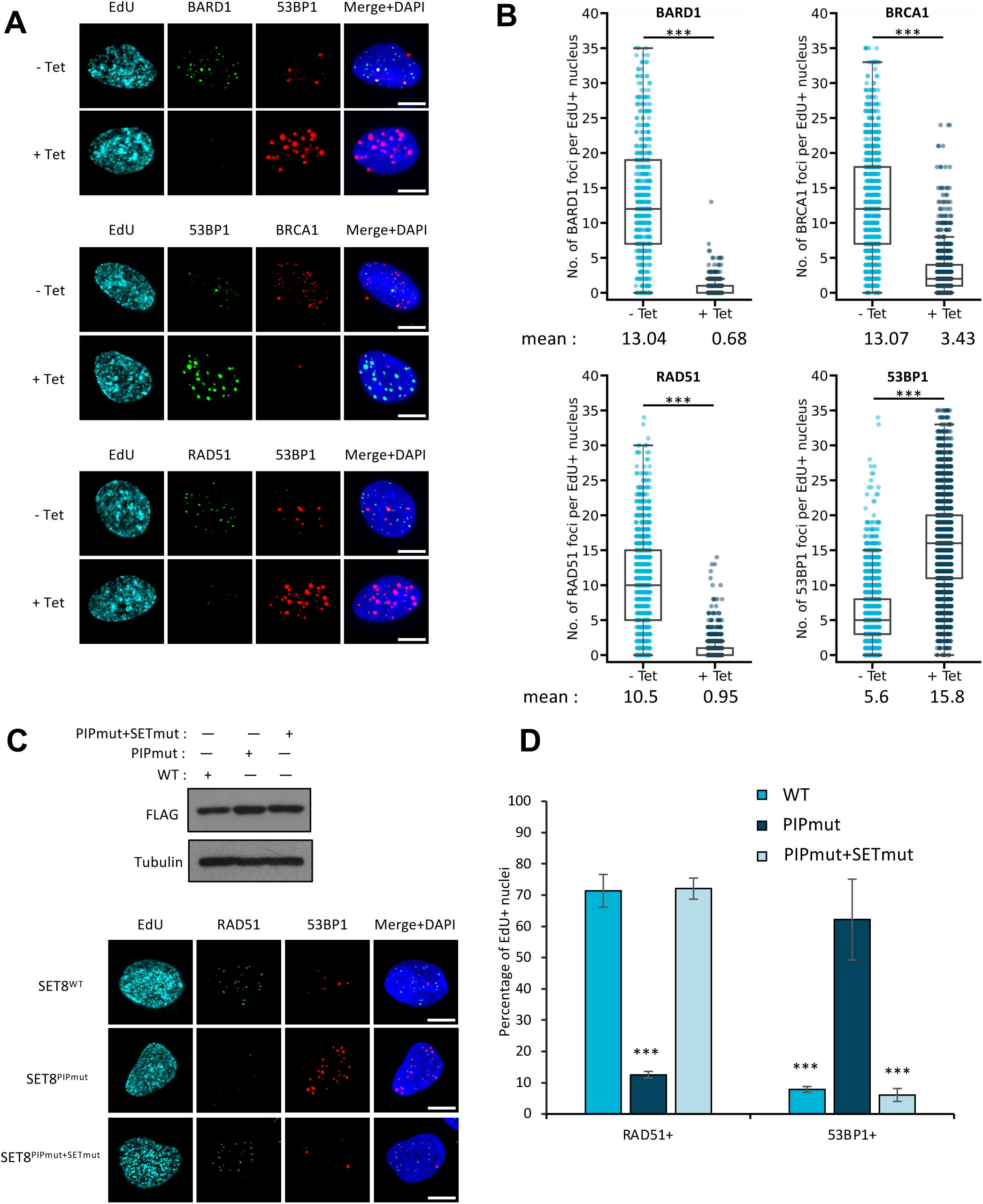
SET8^PIPmut^ expression impairs HR focal accumulation during S-phase. (**A**) Representative images showing staining of 53BP1 with BARD1 (top), BRCA1 (middle), or RAD51 (bottom) in EdU positive nuclei from control (−Tet) and SET8^PIPmut^ (+ Tet). replicating cells. Scale bar = 10 µm. (**B**) Scattered box-plot representing the number of BARD1 (top left), BRCA1 (top right), RAD51 (bottom left) and 53BP1 (bottom right) foci per EdU positive nucleus from control (−Tet) and SET8^PIPmut^ (+ Tet) expressing cells. Inside the box-plot graphs, the thick line represents the median, the limit of the boxes corresponds to the 0.25 and 0.75 quartiles with whiskers showing values 1.5 times the interquartile range. n ≥ 3. ***, p <0.001. Number of nuclei per condition> 100. (**C**). Upper panel: Immunoblots showing similar expression levels of SET8 protein wild-type (WT) and mutants as indicated. Lower panel is representative images showing RAD51 and 53BP1 immunostaining in EdU positive nuclei from replicating cells expressing the different FLAG-SET8 proteins as indicated. (**D**) Bar-plot representing the percentages of EdU positive nuclei that displayed RAD51 or 53BP1 foci in replicating cells expressing the different FLAG-SET8 proteins. Data = mean ± s.d., n ≥ 3. *** p<0.001 (unpaired t-test). Number of nuclei per condition >100.

### The SET8-induced inhibition of HR foci formation depends on 53BP1

Defects in BRCA1 and RAD51 foci formation during DNA replication lead to DNA damage [28–30]. Consistent with this, we noticed larger and higher levels of γH2A.X foci at late time points in SET8^PIPmut^ replicating cells (Figures S1C), which coincided with the activation of the DNA damage checkpoint in G2 cells [23]. To determine whether the recruitment and focal accumulation of 53BP1 upon H4K20me0 depletion is a cause or a consequence of the loss of HR foci formation in replicating cells, we examined the levels of BRCA1, BARD1 and RAD51 foci formation in control and FLAG-SET8^PIPmut^ cells depleted for 53BP1. Two days after two rounds of siRNA transfection to efficiently deplete 53BP1, cells were treated with tetracycline to induce FLAG-SET8^PIPmut^ expression and then pulse-labelled with EdU before to evaluate HR nuclear foci formation during S-phase. Immunoblot experiments verified the efficiency of 53BP1 depletion by siRNA treatment and showed that, independently of 53BP1, FLAG-SET8^PIPmut^ expression triggers the depletion of H4K20me0 recognized by BARD1 (Figure 3A). Accordingly, BARD1 focal accumulation was not properly restored in FLAG-SET8^PIPmut^ replicating cells depleted of 53BP1 (Figure S2A). Consistent with this observation, immunoblot analysis following biochemical fractionation revealed a shift in BARD1 protein levels from the P3 chromatin-enriched fraction to the S1 fraction, which contains soluble cytoplasmic and nuclear components, in control as well as in 53BP1-depleted cells released into S-phase and treated with tetracycline (Figure S2B). Yet, in contrast to BARD1, depletion of 53BP1 was sufficient to fully restore BRCA1 focal accumulation and, to a lesser extent, RAD51 foci formation in FLAG-SET8^PIPmut^ replicating cells (Figures 3B and 3C). These findings suggest that the disruption of BRCA1 and RAD51 nuclear foci formation upon FLAG-SET8^PIPmut^ expression is not necessarily caused by the depletion of H4K20me0 and the loss of its recognition by BARD1, but is more likely dependent on the recruitment and accumulation of 53BP1 mediated by the stabilization of SET8 on post-replicated chromatin.

**Figure 3:**
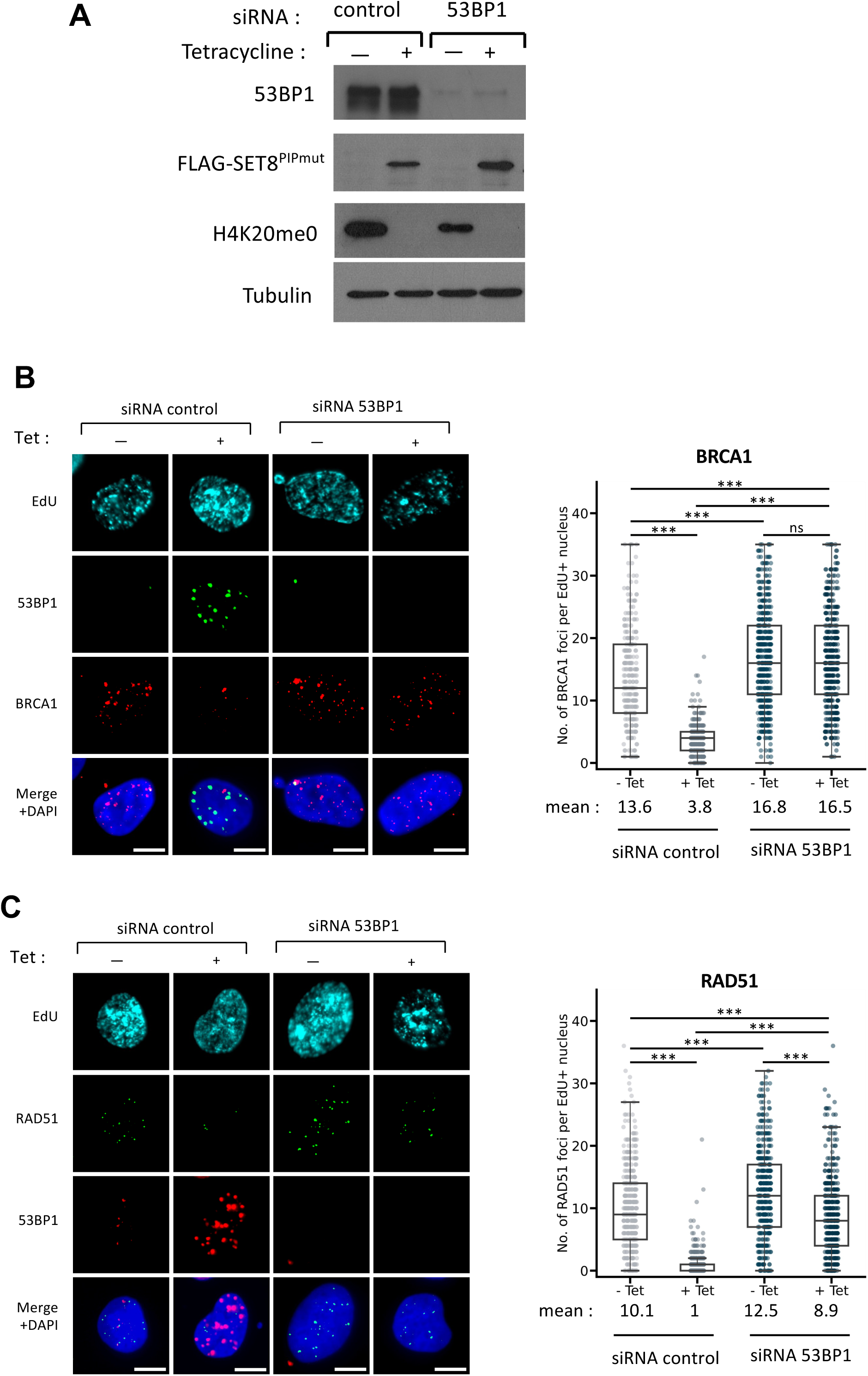
SET8-mediated inhibition of BRCA1 HR functions depends on 53BP1. (**A**) Immunoblots showing the levels of 53BP1, H4K20me0 and FLAG-SET8^PIPmut^ protein in untreated (−Tet) and tetracycline treated FLAG-SET8^PIPmut^ (+Tet) cells and submitted to treatment with control or 53BP1 siRNA as indicated. Tubulin was used as loading control. (**B**) Representative images showing staining of BRCA1 and 53BP1 in EdU positive nuclei from control and 53BP1 siRNA treated cells that were subsequent induced (+ Tet) or not (−Tet) for the expression of the FLAG-SET8^PIPmut^ protein. Scale bar = 10 µm. Right panel is a scattered box-plot representing the number of BRCA1 foci per EdU positive nucleus from the same cells. n=3, *** p<0.001. (**C**). Left panel is representative images showing the staining of RAD51 and 53BP1 in EdU-positive nuclei from control and 53BP1 siRNA treated cells that were subsequent induced (+ Tet) or not (−Tet) for the expression of the FLAG-SET8^PIPmut^ protein. Scale bar = 10 µm. Right panel is the scattered box-plot representing the number of RAD51 foci per EdU positive nucleus from the same cells. Inside the box-plot graphs, the thick line represents the median, the limit of the boxes corresponds to the 0.25 and 0.75 quartiles with whiskers showing values 1.5 times the interquartile range. n=3, *** p<0.001

### The recruitment of 53BP1 on post-replicated chromatin is independent of the subsequent conversion of H4K20me1 to higher H4K20me states

A previous report has proposed that the inverse relationship between 53BP1 and BRCA1 chromatin recruitment is triggered by the replication-coupled dilution of H4K20me2, which is a high-affinity binding site for 53BP1 [9,31]. Hence, the switch from H4K20me0 to H4K20me1 and its subsequent conversion to H4K20me2 should be required for the accumulation of 53BP1 on post-replicated chromatin. To test this hypothesis, the focal accumulation of 53BP1 and HR proteins were examined as described above in Tet-inducible FLAG-SET8^PIPmut^ cells treated or not with tetracycline and the small-molecule inhibitor A196, which specifically targets the family of SUV4-20Hs enzymes responsible for the conversion of H4K20me1 to higher H4K20me states during the cell cycle. The treatment with 10mM of A196 was sufficient to inhibit *de novo* SUV4-20H activity and inhibit the potential conversion of H4K20me1 to higher H4K20me states in FLAG-SET8^PIPmut^ expressing cells (Figure S3). As shown in Figure 4, the focal accumulation of 53BP1 and the concomitant loss of BARD1, BRCA1 and RAD51 nuclear foci during S phase were identical between FLAG-SET8^PIPmut^ expressing cells, whether they were treated or not with A196 (Figure 4). Similar results were obtained with cells treated with A196 for 5 days and that displayed a complete loss of H4K20me2/me3 marks before tetracycline treatment to induce FLAG-SET8^PIPmut^ expression (Figure S4). Taken together, these results show that 53BP1 binding to post-replicated chromatin and the following inhibition of HR focal accumulation are independent of SUV4-20Hs activity and the conversion of H4K20me1 to higher H4K20me states.

**Figure 4:**
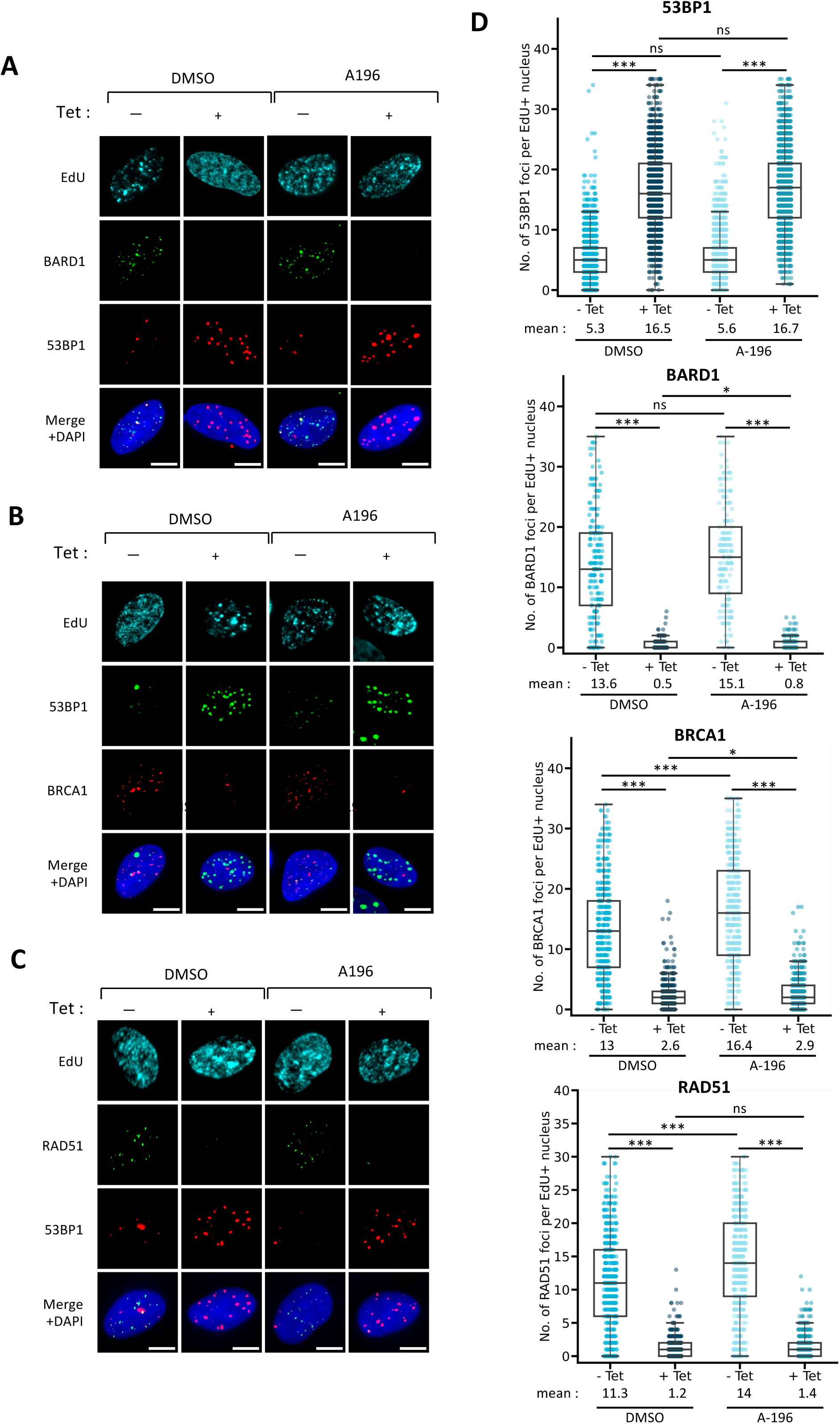
The pharmacological inhibition of SUV4-20H activity did not prevent 53BP1-induced loss of HR nuclear foci upon SET8 stabilization. (**A**) Representative images showing the staining of BARD1 and 53BP1 in EdU-positive nuclei from control (−Tet) and tetracycline (+ Tet) treated inducible FLAG-SET8^PIPmut^ cells, which were previously incubated with DMSO or with 10mM of the SUV4-20H inhibitor A196 for 24 hours. Scale bar = 10 µm. (**B**) Representative images showing staining of BRCA1 and 53BP1 in EdU-positive nuclei from control (Tet) and tetracycline-treated (+ Tet) FLAG-SET8^PIPmut^ cells in presence or DMSO or A196 as indicated in A. Scale bar = 10 µm. (**C**) Representative images showing staining of RAD51 and 53BP1 in EdU-positive nuclei from control (−Tet) and tetracycline-treated (+ Tet) FLAG-SET8^PIPmut^ cells in presence or DMSO or A196 as indicated in A. Scale bar = 10 µm. (**D**) Scattered box-plots representing the number of 53BP1 foci (upper panel), BARD1 foci (middle up), BRCA1 foci (middle down) and RAD51 foci (lowest panel) per EdU positive nucleus from control (−Tet) and FLAG-Set8^PIPmut^ (+ Tet) expressing cells that were incubated with DMSO or A196 as indicated above. Statistical significance was detected when a Student’s test (t-test) was performed with p<0.001. Inside the box-plot graphs, the thick line represents the median, the limit of the boxes corresponds to the 0.25 and 0.75 quartiles with whiskers showing values 1.5 times the interquartile range interquartile range. n=3. Number of nuclei per experiment > 100. * p<0.05; *** p<0.001.

### SET8 cooperates with RNF168 for tipping the balance from BRCA1 to 53BP1 on post-replicated chromatin

By inducing the ubiquitination of histones and others factors, the ubiquitin ligase RNF168 has been reported to be an upstream regulator of 53BP1 that is critical for HR defects caused by BRCA1 deficiency [32–34]. Interestingly, SET8 can interact with RNF168 and promote RNF168 ubiquitin activity towards histone H2A [35,36], thereby raising the possibility that 53BP1 focal accumulation induced by SET8 stabilization could require RNF168. Consistent with this possibility, FLAG-SET8^PIPmut^ expressing cells displayed RNF168 focal accumulation that partially overlapped with 53BP1 (Figures 5A and Figure S5A). This was accompanied by an apparent enrichment of ubiquitinated chromatin proteins as observed by the detection of 53BP1 foci positive for the anti-ubiquitinylated proteins antibody FK2 (Figure 5B). Noted of, the focal accumulation of RNF168 was not reduced upon 53BP1 depletion (Figure S5B) or when SUV4-20H enzymes were inhibited (Figure S5C), thereby indicating that the recruitment of RNF168 on replicated chromatin upon SET8 stabilization occurs independently of 53BP1 and the potential conversion of H4K20me1 to higher H4K20me states. Conversely, to determine whether the recruitment of RNF168 in response to SET8 stabilization contributes to the recruitment of 53BP1 on replicated chromatin and the following inhibition of replication-coupled BRCA1 focal accumulation, we examined by indirect immunofluorescence the levels of 53BP1, BRCA1 and RAD51 nuclear foci in control and FLAG-SET8^PIPmut^ replicating cells previously depleted for RNF168 by siRNA. The depletion of RNF168 and the expression of FLAG-SET8^PIPmut^ were verified by immunoblotting (Figure S5D). As shown in Figures 5C and 5D (left panels), depletion of RNF168 significantly decreased 5BP1 focal accumulation in FLAG-SET8^PIPmut^ replicating cells, thereby suggesting that RNF168 contributes indeed to the untimely accumulation of 53BP1 orchestrated by SET8 stabilization on replicated chromatin. However, although 53BP1 focal accumulation was strongly reduced upon RNF168 depletion, BRCA1 and RAD51 foci formation was not rescued (Figures 5C, 5D and Figure S5E). Instead, we even observed a decrease in HR foci in control cells depleted for RNF168 compared to siRNA control cells visualized by the detection of RAD51 foci (Figure S5E). This was consistent with previous reports showing that RNF168 activity is also required for BRCA1 recruitment, likely via the ability of BARD1 to interact with histone H2A ubiquitination [32,p.168,37]. Altogether, our results suggest a scenario where SET8 stabilization on replicated chromatin subsequently recruits RNF168 and stimulates protein ubiquitination that, in parallel to the single conversion of H4K20me0 to H4K20me1, lead to a post-replicated chromatin organization attractive refractory to BRCA1-dependent HR in a manner depending on the presence of 53BP1.

**Figure 5:**
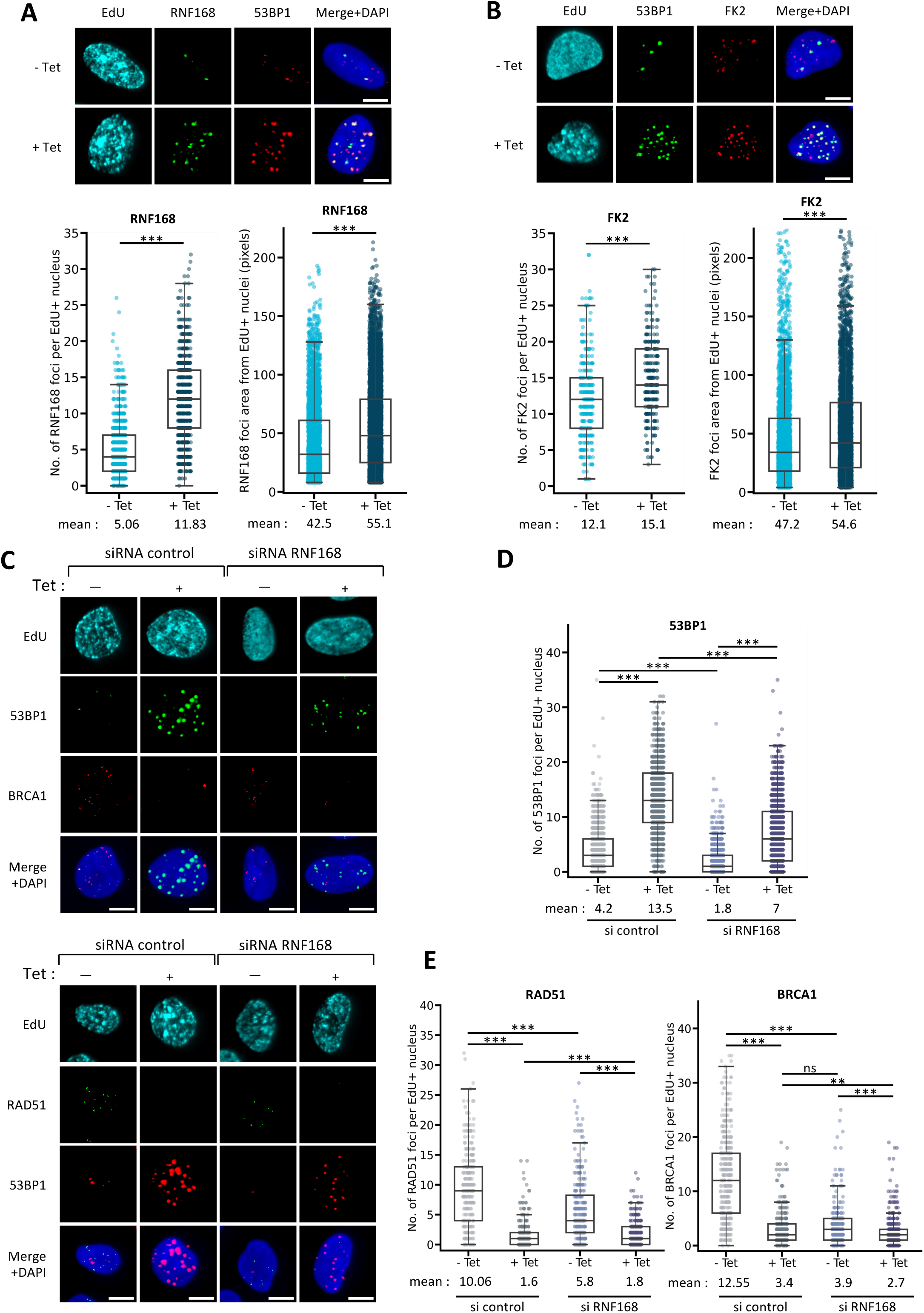
RNF168 and chromatin ubiquitination contribute 53BP1 focal accumulation upon SET8 stabilization. **(A).** Upper panels are representative images of the staining of RNF168 and 53BP1 in EdU-positive nuclei from untreated (−Tet) and tetracycline-treated (+Tet) FLAG-SET8^PIPmut^ cells. Scale bar = 10 µm. Lower panels are scattered box-plots representing the number (left) and area (right) of RNF168 foci in EdU-positive nuclei from untreated (−Tet) and tetracycline-treated (+Tet) FLAG-SET8^PIPmut^ cells. number of nuclei >100 per experiment; n=2. *** p<0.001**. (B).** Upper panels are representative images showing the staining of chromatin ubiquitination (FK2) and 53BP1 in EdU-positive nuclei from untreated (−Tet) and tetracycline-treated (+Tet) FLAG-SET8^PIPmut^ cells. Scale bar = 10 µm. Lower panel are scattered box-plots representing the number (left) and area (right) of chromatin ubiquitination (FK2) foci in EdU-positive nuclei from untreated (−Tet) and tetracycline-treated (+Tet) Tet-inducible FLAG-SET8^PIPmut^ cells. Number of analyzed nuclei >100 in this experiment; *** p<0.001. **(C)** Representative images of the staining of 53BP1, BRCA1 and RAD51 in EdU-positive nuclei from untreated (−Tet) and tetracycline-treated (+Tet) Tet-inducible FLAG-SET8^PIPmut^ cells that were transfected with control or RNF168 siRNA two days before tetracycline treatment. Scale bar = 10 µm. n=2. **(D)** Scattered box-plot representing the number of 53PB1 foci in the nuclei as indicated in C. n=2, Number of nuclei >100 per experiment. *** p<0.001. (**E)** scattered box-plot representing the number of RAD51 and BRCA1 foci in the nuclei as indicated in C. number of nuclei >100 per experiment. ** p<0.01; *** p<0.001.

### SET8 is a potent cell-cycle regulated inhibitor of homologous recombination

The findings described above on the inhibitory role of SET8 in replication-coupled HR foci formation suggest that this epigenetic enzyme functions as a chromatin inhibitor of homologous recombination. To explore further this possibility, control (−TET) and SET8^PIPmut^ expressing cells (+TET) were treated with camptothecin (CPT) to induce DNA damage and then the number of RAD51, BRCA1, 53BP1 and RNF168 nuclear foci were examined by indirect immunofluorescence. Camptothecin (CPT) is a Topoisomerase-I poison that promotes DNA breakage that are mainly repaired by HR in U2OS cells [38]. Accordingly, CPT led to the activation of the DNA damage checkpoint kinase ATM (Figure S6A) followed by the focal accumulation of RAD51, BRCA1 and 53PB1 in a dose-dependent manner in control cells (Figures 6A and 6B). In contrast, although ATM is still normally activated in tetracycline-treated cells upon CPT treatment (Figure S6A), FLAG-SET8^PIPmut^ expression inhibited the focal accumulation of RAD51 and BRCA1, but not of 53BP1 that was even increased (Figures 6A and 6B). The focal accumulation of RNF168 was also enhanced (Figure S6B). These results were consistent with a previous report showing that the overexpression of SET8 reduces the loading of BRCA1 and RAD51 on chromatin upon ionizing radiation [31]. Importantly, RPA foci formation on DNA lesions was observed in both control and SET8^PIPmut^ expressing cells upon CPT treatment (Figure S6C), indicating that the loss of RAD51 foci formation was not indirectly caused by defects in RPA loading at DNA breaks or defects in DNA end resection. Consistent with the accumulation of 53BP1 instead of BRCA1, we also observed the specific phosphorylation of DNA-PKcs at Serine 2056 upon CPT in FLAG-SET8^PIPmut^ expressing cells (Figure S6A), thereby suggesting the activation of NHEJ pathway rather than the HR pathway. Finally, upon 53BP1 depletion and consistent with the results shown in Figures 3A and 3B, the number of RAD51 and BRCA1 foci was largely restored in FLAG-SET8^PIPmut^ replicating cells treated with CPT, thereby confirming the requirement of 53BP1 in the SET8-mediated inhibition of HR foci (Figure S6D).

**Figure 6:**
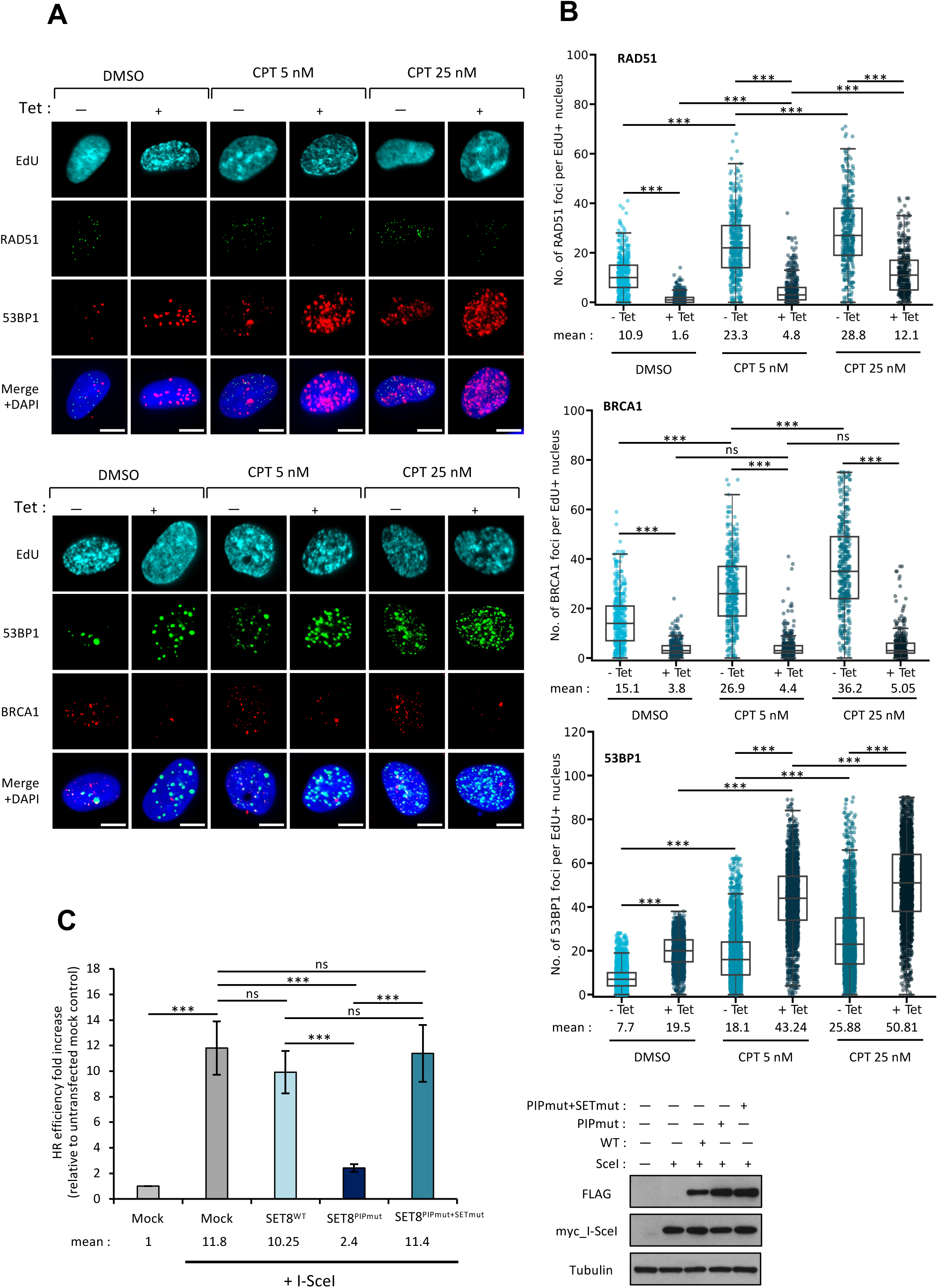
SET8 inhibits homologous recombination. (**A**) Representative images of RAD51, BRCA1 and 53BP1 staining in EdU positive nuclei from control (−Tet) and FLAG-SET8^PIPmut^ (+Tet) expressing cells treated with 5 or 25 nM of camptothecin (CPT) during 2 hours or with vehicle (DMSO) as indicated. Scale bar = 10 µm. (**B**) Scattered box-plot representing the quantification of RAD51, BRCA1 and 53BP1 foci per EdU-positive nuclei from control (−Tet) and FLAG-SET8^PIPmut^ expressing cells (+ Tet) treated with 5 nM or 25 nM of camptothecin during 2 hours, or with vehicle (DMSO). Interquartile range and statistical significance are shown as in Figure 2A. n ≥ 3; *** p<0.001 (t test). Number of nuclei per condition and experiment >100. (**C**) Left panel: Bar-plot representing the relative efficiency of DNA repair by homologous recombination of I-SceI-induced DNA breaks as evaluated by flow cytometry using the U2OS DR-GFP cell line after expression of different FLAG-SET8 proteins as indicated. Data = mean ± s.d., n = 3. *** p<0.001 (t-test). Number of events per sample > 10 000. Right panel: immunoblots showing the similar expression of different FLAG-SET8 proteins and of MYC-tagged nuclease I-SceI in U2OS DR-GFP cells. Tubulin was used as loading control.

To corroborate further the inhibitory role of SET8 in homologous recombination, the efficiency of HR upon SET8 expression was assessed using a DR-GFP reporter-based HR system stably integrated into the genome of U2OS cells. This DR-GFP system consists of two unique inactive copies of the GFP gene integrated at the same loci. The first GFP copy (SceGFP) contains the I-SceI restriction site with an in-frame stop codon and the second copy (secGFP) harbors truncations at both ends. After cleavage within SceGFP copy by I-SceI, mostly HR uses secGFP as a template to restore a functional GFP gene. Then, GFP fluorescence can be detected using flow cytometry as an indirect measure of HR efficiency. DR-GFP cells were first transduced with retroviral vectors encoding either FLAG-SET8^WT^, FLAG-SET8^PIPmut^ or the inactive FLAG-SET8^PIPmut+SETmut^ protein and then transfected with a plasmid expressing Sce1 (Figure 6C). Consistent with SET8-induced defects in HR foci formation in presence of exogenous DNA damage (Figures 6A and 6B), the expression of FLAG-SET8^PIPmut^ strongly reduced the efficiency of HR in a manner depending on its methyltransferase activity (figure 6C). As expected, however, the HR efficiency was only slightly reduced in cells expressing FLAG-SET8^WT^ (Figure 6C), which was still highly degraded during S and G2 phases when HR is active [23]. Taken together, these results demonstrate that SET8 works as a potent inhibitor of homologous recombination and that the proper activation of this DNA repair pathway during S/G2 phases requires the PCNA-coupled degradation of SET8 in order to avoid the inhibition of BRCA1 signaling functions in recombination mechanisms.

### SET8 contributes to curtail BRCA1 chromatin binding outside of S/G2 phases

In order to avoid genetic aberrations, the functions of BRCA1 in homologous recombination must be suppressed from G1 phase to early S-phase, when only homologous chromosomes are available [3]. Based on the findings described above and that chromatin is enriched in H4K20me1 during G1 and early-S phases, we reasoned that SET8 methyltransferase activity on histone H4K20 might contribute to limit the activity of BRCA1 outside of S/G2 phases. Consistent with this hypothesis, it was previously shown that SET8 depleted cells display higher BRCA1 chromatin binding [12]. However, it was uncleared whether this was due to the lack of conversion of H4K20me0 to H4K20me1 states after DNA replication. To address this issue, U2OS cells were synchronized at G1/S transition by double thymidine block, transfected with control or SET8 siRNA for 12 hours, then released from the G1/S transition before FACS analysis of cell-cycle progression. Both control and SET8 siRNA-treated cells were able to progress normally and similarly through S-phase, then exited from mitosis 15 hours after release with, however, high levels of H4K20me0 instead of H4K20me1 in SET8 depleted cells (Supplementary Figure S7). At 15 hours post release and compared to control cells, immunoblots after biochemical fractionation showed the loss of BRCA1 in S1 fraction of cytoplasmic components and its concomitant enrichment in P3 chromatin-enriched fraction of SET8 depleted cells harboring H4K20me0. In contrast, the chromatin binding of 53BP1 was slightly but significantly decreased, as observed by the lower levels of 53BP1 in P3 fraction and its concomitant increase in S2 fraction with soluble nuclear components (Figure 7A). In a previous study, we showed that G1/S synchronized cells depleted for SET8 started to accumulate DNA breaks as they progressed from G1 to S phase [39]. In agreement with this, FACS analysis showed that synchronized control cells continued to progress through the cell cycle, whereas an accumulation at G1/S transition was observed for SET8 depleted cells (Figures 7A and 7B). EdU pulse labelling followed by immunofluorescence showed that this G1/S arrest was accompanied by a focal accumulation for BRCA1, but not for 53BP1, in most of SET8 depleted cells (Figure 7C). In contrast, control G1 cells negative for EdU only displayed rare BRCA1 foci, which emerged when control cells progressed into S phase as expected (Figure 7C). Altogether, these results showed that the loss of SET8, and likely the absence of conversion of H4K20me0 to H4K20me1 after DNA replication, lead to a decrease in 53BP1 chromatin binding at the subsequent cell cycle. This decrease is associated with improper focal accumulation of BRCA1 in chromatin and defects in the G1 to S phase transition.

**Figure 7:**
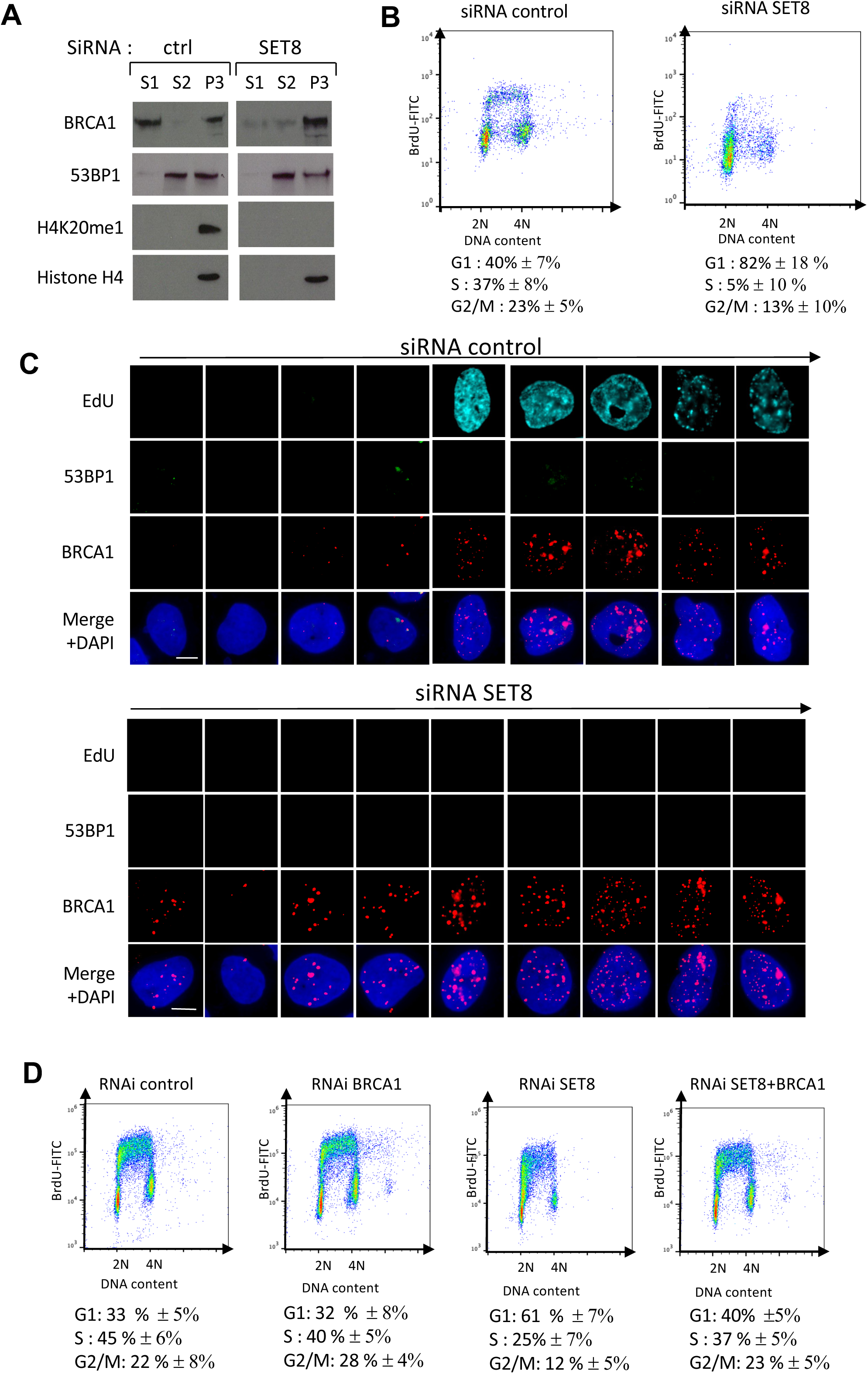
SET8 depletion induces improper chromatin and focal accumulation of BRCA1 outside of S-phase. (**A)**. Representative immunoblots showing the localization of BRCA1 and 53BP1 in soluble cytoplasmic (S1) and nuclei (S2) fractions and in chromatin-enriched fractions (P3) at early G1 phase in control and SET8 SiRNA-treated cells that excited from mitosis. Cells were harvested 15 hours after G1/S release. n=3 (**B**) FACS analysis of DNA content and BrdU signal after mitotic exit of the control and SET8 SiRNA-treated cells as described above. Cells were harvested 24 hours after G1/S release. n=3 (**C**) Representative images showing EdU incorporation (DNA synthesis) and staining of 53BP1 and BRCA1 in control and siRNA SET8 treated cells progressing from G1 to S-phase after a first mitosis without SET8 and H4K20me1 as described above. Cell synchronization was performed as depicted in Supplementary Figure S7. Scale bar = 10 µm (**D**). FACS analysis of DNA content and BrdU incorporation levels of control (shLuc/siCtrl), SET8-depleted U2OS cells (shRNA SET8/siRNA Ctrl), BRCA1-depleted U2OS cells (shRNA Luc/siRNA BRCA1) and cells depleted for both BRCA1 and SET8 (shRNA SET8/siRNA BRCA1). n=3. Number of events per sample > 10 000.

### Improper BRCA1 chromatin accumulation contributes to G1-S progression defects in SET8 depleted cells

Since the inaccurate BRCA1 foci formation outside of S-phase could have deleterious cellular effects [40], we wondered whether BRCA1 could contribute to the accumulation of SET8 depleted cells at G1/S transition. To address this question, U2OS cells were first transduced with retroviral vectors expressing shRNA control or shRNA SET8 and the following days transduced cells were transfected with a control siRNA or a siRNA directed against BRCA1. Two days after treatment, cells were pulse-labeled with BrdU and DNA replication progression were analyzed by FACS. Whereas BRCA1 siRNA treatment by its own did not affect S phase entry as measured by FACS analysis, depletion of BRCA1 was sufficient to partially restore BrdU incorporation and normal S-phase progression in SET8 depleted cells (Figure 7D). These results suggest an inaccurate binding of BRCA1 on chromatin participate to the G1/S arrest upon SET8 depletion. Taken together, our results demonstrate the importance of the timely *de novo* SET8-induced H4K20 mono-methylation in the regulatory cell-cycle inhibition of BRCA1 signaling in absence of exogenous source of DNA damage, an essential mechanism for the maintenance of genome integrity during normal cell proliferation.

**Figure 8:**
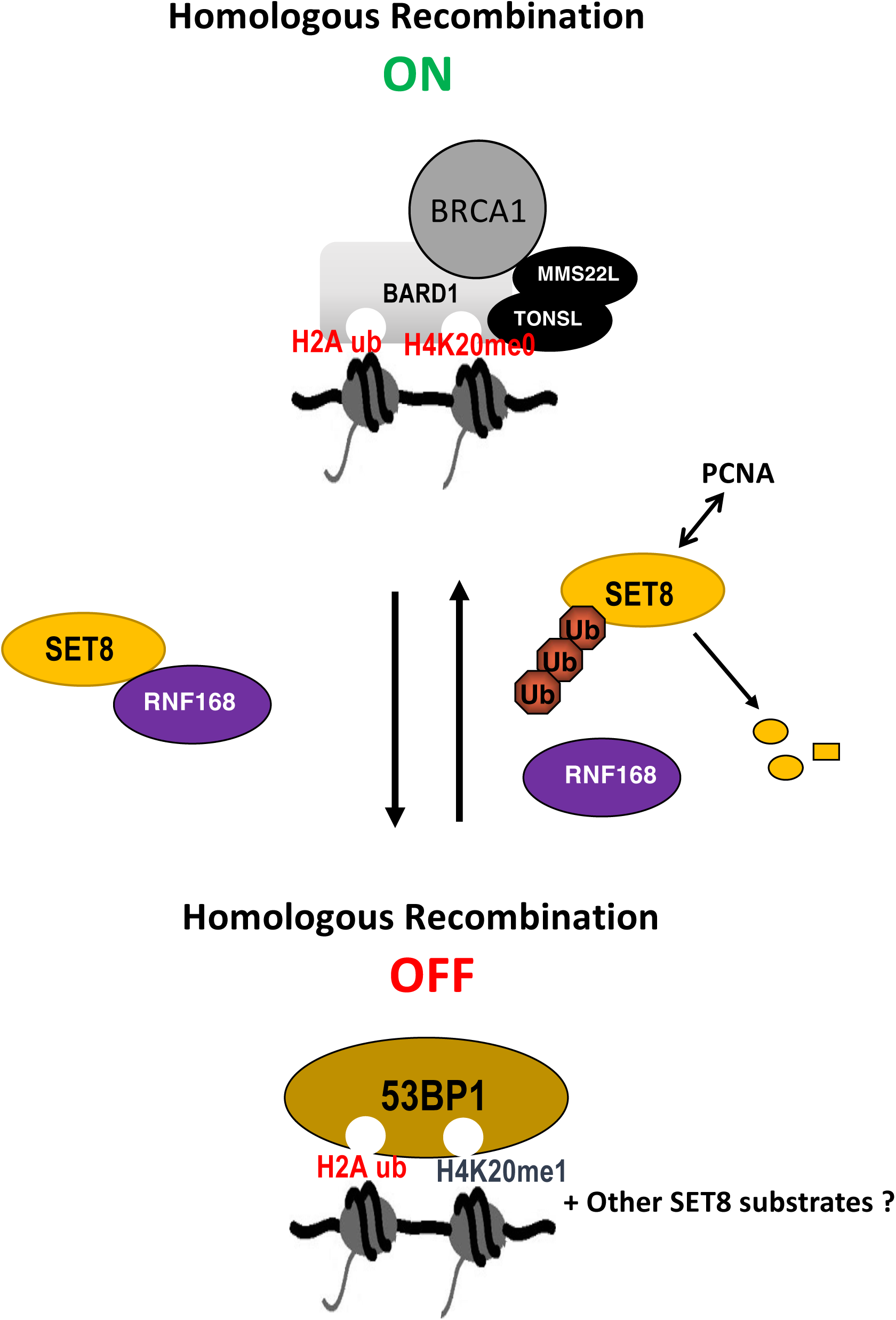
Proposed model for the cell cycle-mediated regulation of BRCA1 and 53BP1 chromatin accumulation by the concerted activities of SET8 and RNF168 in absence of exogenous source of DNA damage. The concerted activities of the H4K20 mono- methyltransferase SET8 and the ubiquitin ligase RNF168 responsible for histone H2A ubiquitination serve as turn-off on system for homologous recombination pathway by tipping the balance from BRCA1 to 53BP1 complexes during the cell cycle. This likely depends on the conversion of unmethylated H4K20 (H4K20me0) to monomethylated H4K20 (H4K20me1), without excluding the possibility of other SET8 substrates that remain to be identified. The proteolytic regulation of SET8 during the cell cycle determines when and where occurs the switch from BRCA1 to 53BP1 complexes. This occurs independently of the subsequent conversion of H4K20me1 to higher H4K20 methylation states by SUV4-20H enzymes.

## DISCUSSION

While our knowledge of the mechanisms that activate BRCA1-mediated HR pathways upon DNA damage is extensive, the mechanisms that regulate BRCA1 activity upon unchallenged cell-cycle conditions remains poorly understood. Yet, BRCA1-mediated HR signaling is required for repair of spontaneous double-stranded breaks (DSBs) that may arise during DNA replication, a function that must be tightly controlled to ensure proper cell cycle progression and avoid pathological situations such as in breast or ovarian carcinomas [8]. In this regard, we provide new evidence on the role of the replication-coupled degradation of the lysine methyltransferase SET8 responsible for H4K20 mono-methylation (H4K20me1) and demonstrates that this proteolytic regulation is primordial for the onset of the recombinogenic functions of BRCA1 during DNA replication. In cooperation with the ubiquitin ligase activity of RNF168, we show that the methyltransferase activity of SET8 on post-replicated chromatin is required to turn off homologous recombination by tipping the balance from BRCA1-BARD1 to 53BP1 complexes. This is likely related to the switch from un-methyl to mono-methyl K20 on newly incorporated histones H4, although we cannot exclude that other SET8 non histones substrates, such as P53 or PCNA [41,42], could also contribute to balance between BRCA1 and 53BP1 activities. Conversely, the lack of SET8 and restoration of H4K20me1 marks after DNA replication led to the chromatin accumulation of BRCA1 instead of 53BP1 in the subsequent G1 phase, thereby causing chromatin and replication defects in daughter cells. Taken together and as illustrated in the proposed model in Figure 7, these results clarify the mechanisms by which SET8 could impact on DNA repair pathway choice. Specifically, this establishes the *de novo* activity of SET8 on post-replicated chromatin as the primordial epigenetic lock of BRCA1-mediated HR pathway during the cell cycle in unperturbed cells.

The results described here are consistent with a previous study showing that the overexpression of SET8 reduces BRCA1 and RAD51 foci formation and inversely promotes 53BP1 focal accumulation on damaged chromatin upon ionizing radiation [31]. However, in this previous study, the conversion of H4K20me1 to H4K20me2 appeared to function downstream of SET8 in 53BP1 chromatin recruitment at DNA breaks [31]. Our results do not favor this hypothesis, where the recruitment of 53BP1 on post-replicative chromatin appear independent from the activity of the enzymes responsible for the conversion of H4K20me1 to higher H4K20me states. Strikingly, several studies have reported conflicting results regarding the H4K20me states and the H4K20 enzymes that dictate the recruitment of 53BP1 on damaged chromatin. Whereas the decrease in H4K20me2/me3 upon the SUV4-20H inhibitor A196 reduces 53BP1 foci formation upon ionizing radiation in cancer cells [43], the loss of SUV4-20H enzymes and the absence of H4K20me2/3 only induced a slight delay on 53BP1 focal accumulation at DNA breaks in mouse embryonic fibroblasts [44]. We believe that this apparent discrepancy ultimately reflects the disparate *in vivo* affinity of 53BP1 for the different H4K20me states. This likely translates into different modes of 53BP1 recruitment depending on the level and nature of DNA damage sustained, the chromatin epigenetic landscape, and the phases of the cell cycle. Thus, our results demonstrate that the 53BP1 recruitment to chromatin in the absence of exogenous DNA damage is linked to SET8’s dual functions: (i) the ability of SET8-induced H4K20me1 to evict H4K20me0 readers and their associated HR factors notably BRCA1 and (ii) the ability of SET8 to create efficient 53BP1 binding sites along the genome via the switch from H4K20me0 to H4K20me1 and by promoting RNF168-mediated chromatin ubiquitination.

A potential mechanism by which SET8 could stimulate RNF168 activity is suggested by two recent studies showing that SET8 directly interacts with RNF168 and that this interaction is important for SET8 recruitment to DNA damage while enhancing RNF168 ubiquitin activity on chromatin proteins [35,36]. In this context, it will be valuable to investigate how non-degradable SET8 mutants in combination with other mutations, such as a SET8 enzyme unable to interact with RNF168, affect histone ubiquitination and the DNA-repair activity of associated readers 53BP1 and BARD1 on post-replicated chromatin. Additionally, it will worth it to examine the impact of SET8 mutants on the enzymes that regulate RNF168 activity on chromatin, such as the E3 ubiquitin ligases TRIP12 and UBR5 that are normally recruited to replicated chromatin in order to limit histone ubiquitination [45].

It is well established that HR defects in BRCA1-deficient cells can be compensated by the inactivation of 53BP1 [45]. Furthermore, depending on the mechanism of BRCA1 inactivation, 53BP1 inhibits either DNA resection step or the deposition of RAD51 on DNA post-resection [40]. Since H4K20me0 is a post-replicative mark during S-phase and is incompatible with 53BP1 recruitment, it has been suggested that H4K20me0 recognition by BARD1 tips the balance of DNA repair in favor of BRCA1 and HR factors [12,17]. However, our data do not fully support this model. Indeed, we show that the depletion of 53BP1 is sufficient to restore BRCA1 and RAD51 focal accumulation even in absence of H4K20me0 and proper levels of BARD1 foci formation (Figure 3). Therefore, our results suggest that the recruitment of BRCA1 on replicated chromatin does not necessary depend on the recognition of H4K20me0 by BARD1. Given that BARD1 can function independently of BRCA1 [46], it remains to determine whether H4K20me0 levels and the activity of SET8 might play a critical role in BRCA1-independent functions of BARD1. Of particular interest would be investigating how the methylation of H4K20me0 influences BARD1 role in p53-mediated apoptotic responses, as SET8 can also target p53 transcriptional functions by inducing the mono-methylation of lysine 382 [41]. An intriguing question remains: How BRCA1 focal accumulation can be restored without H4K20me0 and BARD1? This could be related to the ability of BRCA1 to form distinct protein complexes through the association of different partners [47]. Another possibility, though not mutually exclusive, is that SET8-induced recruitment of RNF168 contributes to restore nuclear BRCA1 and RAD51 foci in absence of 53BP1. Indeed, recent findings show that RNF168 interacts with PALB2, which facilitates RAD51 loading onto DNA when BRCA1 homologous recombination functions are impaired [33].

Another significative finding of our study is that the absence of SET8 and of the restoration of H4K20me1 signal after DNA replication leads to the excessive chromatin binding and focal accumulation of BRCA1 (Figure 7). In contrast, we observed the reverse effect on 53BP1 (Figure 7). Previous research has established that the loss of SET8-dependent H4K20 methylation at the end of S-phase led to an immature chromatin organization in the subsequent G1 phase [39]. This chromatin disorganization leads to the accumulation of DNA damage as cells progress into S-phase, primarily due to uncontrolled chromatin loading of the ORC/MCM complex, origin firing and replication stress [39]. We propose that the untimely BRCA1 focal accumulation in pre-replicative cells is related to these chromatin alterations. Although BRCA1 is essential for managing replication-associated DNA damage, our results suggest that its chromatin accumulation before or too early during S-phase may be incompatible with DNA replication initiation. Additionally, our findings suggest that a chromatin with reduced H4K20me1 levels impairs the ability of 53BP1 to bind near DNA lesions, which indirectly creates a chromatin environment more permissive to BRCA1 binding and thus contributes to defective G1 to S phase progression. This is reminiscent of a previous result showing that targeting RAD51 by siRNA partially restored S-phase progression in asynchronous SET8 depleted cells, suggesting an inaccurate activation of HR process in these cells [48]. However, Despite BRCA1 foci formation, we did not observe a significant accumulation of RAD51 foci in SET8 depleted cells arrested at G1/S transition. The exact role played by BRCA1 and RAD51 in cell-cycle defects upon SET8 depletion remains to be determined.

The formation of BRCA1 foci outside of S/G2 phase is not unprecedented, as 53BP1 depletion has been shown to result in ectopic BRCA1 foci following ionizing radiation in G1 cells [49,50]. Interestingly, BRCA1 has also been shown to be naturally recruited at damaged centromeric regions in G1 cells in a manner depending on the histone variant CENP-A and histone H3K4 di-methylation [51], suggesting that the chromatin recruitment of BRCA1 can occur at specialized chromatin regions of the genome independently of DNA replication. While further experiments are necessary to fully understand these mechanisms, it is tempting to imagine that localized control of SET8 activity and the regulation of 53BP1’s interaction with H4K20me along the genome could allow BRCA1 to bind specifically to centromeric regions, while preventing its spread beyond centromeric chromatin during G1 phase.

Clinically, SET8 has been found upregulated in several types of cancers and often associated with poor prognosis [52–54]. The higher-level of SET8 in cancer cells could be due to multiple mechanisms, including DNA copy-number gains, changes in gene expression or enhanced protein stability [55–57]. To date, however, the benefit of high levels of SET8 on cancer cell development and resistance is far from understood. Given the importance of BRCA1 deficiency in tumorigenesis and our novel findings regarding the inhibitory role of SET8 on BRCA1 during the cell cycle, it will be interesting in near future to explore whether SET8 might contribute to the reduction of functional BRCA1 during tumorigenesis. Notably, this issue seems particularly relevant in tumors where BRCA1 dysfunction cannot be easily attributed to mutations or loss of expression, as observed in certain subtypes of sporadic breast and ovarian cancers, where SET8 is often upregulated. In this context, targeting SET8 with inhibitors to restore BRCA1 function represents a promising avenue for further research. However, this will necessitate the development of more selective small-molecule inhibitors with better pharmacokinetic and pharmacodynamic properties than the SET8 inhibitors currently available.

## EXPERIMENTAL PROCEDURES

### Cell culture, protein mutations and chemical drugs

HEK293T, U2OS, DR-GFP, U2OS Tet-On cell lines were cultured at 37°C in Dulbecco’s Modified Eagle Medium (DMEM) with Glutamax (Gibco) and supplemented with 10% Fetal Bovine Serum (FCS), 1% antibiotics (Gibco) and 1% Sodium Pyruvate (Gibco). Expression of the different FLAG-SET8 proteins in U2OS cells were induced by the addition of 2 µg/ml of tetracyclin (TET) (Sigma-Aldrich) into cell culture medium. G1/S synchronization was performed by double thymidine block as described previously. Alternatively, confluent cells arrested in G1 were seeded at low density in presence of 2.5 mM Thymidine (Sigma-Aldrich) for 24 hours. Release of cells from G1/S block was performed by three cycles of washes with PBS then fresh medium for 5 minutes at 37°C. The SUV4-20H inhibitor A196 (Sigma-Aldrich) was used at the concentration of 10 µM. Camptothecin (C9911, Sigma-Aldrich) was used to induce replicative stress and DNA breaks during S-phase. The SET8^PIPmut^ (F184A and Y185A mutations) and SET8^PIP-Setmut^ ^(^F184A, Y185A and D338A mutations) have been described previously [23].

### RNA interference experiments

One day after seeding 10^6^ U2OS cells in 6-well plate, small interfering RNAs (siRNA, 20mM) were delivered using Lipofectamine® RNAi-MAX Reagent (Invitrogen) according to the supplier’s instructions. Briefly, transfection mixture containing 300 µl Optimem (Gibco), siRNA and Lipofectamine® RNAi-MAX Reagent was added to each well containing 1 ml of culture medium for 6 to 8 hours at 37°C before washing the cells with PBS and culturing them in fresh medium. SET8 siRNA sequence was provided and validated by Cell Signaling (#1307). The other validated siRNA sequences were provided by Sigma-Aldrich (MISSION® esiRNA): Control (EHURLUC, EHUFLUC, EHUGFP), TP53BP1 (EHU156121), RAD51 (EHU045521), BRCA1 (EHU096311), RNF168 (EHU011891). Retroviral pSIREN vectors expressing shRNA sequences directed against SET8 or Luciferase (Control) were described previously [58]. Briefly, retroviral particles were collected two days after two rounds (at 24h interval) of plasmid transfection into HEK293T cells. After filtration, retroviral particles were directly added to U2OS cells in the presence of polybrene (8 μg/ml, Sigma-Aldrich). Twenty-four hours after infection, shRNA expressing cells were selected with puromycin (1 mg/ml).

### Cell-Cycle analysis by Flow Cytometry

Cells were incubated with BrdU (50 µg/ml, Sigma-Aldrich) for 90 minutes before fixation in ice-cold 70% ethanol. After incubation for 30 minutes with solution containing 2N HCl and 0.5% Triton X-100 under gentle agitation, cells were washed with Borax buffer (pH8.5) and then sequentially incubated in blocking solution (PBS, 0,5% Tween 20,1% BSA) containing anti-BrdU primary antibody (1/100, BD Biosciences) and anti-mouse secondary antibody coupled to FITC (1/300, Sigma-Aldrich) for 2 hours each. Total DNA was labeled with 2 μg/ml of 7-AAD (7-amino-actinomycin D, Sigma-Aldrich) diluted in PBS containing 100 μg/ml of RNAse A (Sigma-Aldrich). FACS analysis was carried out with Gallios flow cytometer (Beckman Coulter) using the Kaluza Acquisition software (Beckman Coulter). Cell cycle profiles were analyzed using FlowJo software (BD Biosciences). the data were presented with mean ± s.e.m. from at least three independent experiments. Statistical significance was evaluated by an unpaired t-test. ns : p-value > 0.05; * : p-value < 0.05; ** p-value < 0.01; *** : p-value < 0.001.

### Homologous recombination (HR) assays

The HR assays were performed using the U2OS DR-GFP cell line (ATCC) that allows to monitor homologous recombination efficiency through the detection of GFP positive cells via flow cytometry. One day after infection with retroviral empty pBABE vector or expressing FLAG-SET8^WT^, FLAG-SET8^PIPmut^ or FLAG-SET8^PIPmut+SETmut^, cells were transfected with 5 ug of plasmid encoding the I-SceI endonuclease using JetPEI reagent and following manufacturer’s instructions. Cells were then collected two days later, fixed in 1% Formaldehyde for 15 minutes and then permeabilized for 1h at 4 °C. After one hour incubation with 2 µg/ml of 7-AAD, the measure of DNA content and the percentage of GFP positive cells were carried out with Gallios flow cytometer (Beckman Coulter) using the Kaluza Acquisition software (Beckman Coulter). FACS data were analyzed using FlowJo software (BD Biosciences). The results are presented as mean ± s.e.m. from four independent experiments with *p*-values determined with unpaired t-test.

### Immunofluorescence and Statistical analysis

After incubation for 20 minutes with 10 µM of EdU (Invitrogen), cells on microscope slides were treated with ice-cold PBS containing 0.25% Triton for 2 min and then fixed with 4% formaldehyde for 12 min. After permeabilization with PBS containing 0.25% Triton for 10 min, incorporated EdU was chemically conjugated to Alexa 647 fluorochrome using Click-iT Plus EdU Cell Proliferation Kit (Invitrogen). After PBS washing, samples were incubated for one hour in a blocking solution (PBS, 0.1% Tween 20, 5%BSA), before adding the primary antibody for an overnight incubation in a humid chamber at 4°C. After washing in PBS containing 0,1% Tween 20, cells were incubated with secondary antibodies for 2 hours at RT in a humid chamber and DNA was counterstained with 0.1 µg/ml of DAPI (DNA intercalator, 4’,6-Diamidino-2-Phenylindole, D1306, Invitrogen). Coverslips were mounted with ProLong Diamond Antifade (Invitrogen). The following primary antibodies used were anti RPA70 (1:100, #2267, Cell Signaling), anti-53PB1 (1:500, Cell Signaling and 1:1000, Millipore), anti-RNF168 (1:100, GTX129617, Genetex), anti-RAD51 (1:500, PC130, Millipore), anti-gH2A.X (1:1000, Millipore), anti-FK2 (&:250, Millipore), anti-BRCA1 (1:500, sc-6954, Santa-cruz), anti BARD1 (1:500, PLA0074, Sigma). The secondary antibodies used were anti-Rabbit-Alexa Fluor 488 (1/500, Thermo Fisher Scientific) and anti-Mouse-Cy3 (1/500, Thermo Fisher Scientific). Edu staining was revealed with a secondary Alex647 antibody. Slides were analyzed using Zeiss Axio Imager M2 upright microscope (Zeiss) equipped with an Apochromat 63X objective NA 1.4, (immersion oil). Images were strictly acquired under the same microscopic conditions using an ORCA-Flash4.0 LT+ Digital CMOS camera (C1140-42U30, Hamamatsu) controlled by Zen acquisition software (Zeiss). Images were prepared using Zen software and nuclear foci from each image were automatically counted using CellProfiler software. Fiji/ImageJ from the NIH (http://rsbweb.nih.gov/ij/) was used to convert the ZEN format files generated by the Zeiss software into TIFF files recognized by the Cell Profiler software. The analysis was automated by the development of foci counting pipelines (available on request), allowing the automatic counting of nuclear foci from large numbers of images. Values from at least 100 cells per experiment were compiled. Replicating Nuclei were defined using the EdU and DAPI signal. Then, the number, area and intensities of detected foci in replicating nuclei were recorded for the other fluorophores (Alex 488 and Cy3) in database. Quantitative and statistical analyses of database were performed using Python and Prism 9.0 software. Statistical significance was evaluated by unpaired t-test. ns : p-value > 0.05; * : p-value < 0.05; ** p-value < 0.01; *** : p-value < 0.001.

### Protein extraction, immunoblot analysis and small-scale fractionation

Cells were lysed at 95°C for 10 minutes in 2X SDS Sample Buffer (100 mM Tris-HCl pH6.8, 20% Glycerol, 2% SDS 2%, 1 mM DTT, 5% β-mercaptoethanol). Proteins samples were separated on polyacrylamide gel under denaturing conditions (SDS-PAGE) and transferred to nitrocellulose or PVDF membranes following the primary antibodies used. Before incubation with primary and secondary antibodies, membranes were saturated in a solution of TBS containing 0.2% Tween 20 and 5% w/v of dried milk for 16 hours at 4°C. For immunoblots analysis the following antibodies were used : anti-H4K20me0 (1:1000, Abcam), anti-histone H3 (1:5000, Cell Signaling), anti-histone H4 (1:1000, Cell Signaling), anti-H4K20me states (1:1000, cell Signaling), anti-SET8 1:1000, Cell Signaling), anti-53BP1 (1:1000, cell Signaling), anti-BRCA1 (1:1000, cell Signaling), anti-ATM and anti-pS1981-ATM (1:1000, cell Signaling), anti-DNA-PK unphosphorylated (1:1000, Millipore) and phosphorylated at S2056 (1:1000, Cell Signaling), anti-RNF168 (1:1000 Genetex), anti-FLAG (1:1000, Sigma) and anti-Tubulin (1:1000, Sigma). The following secondary antibodies were used anti-rabbit (1/10000, Cell Signaling) and anti-mouse (1 /10000, Cell Signaling). Protein bands were visualized on X-ray film by electro-chemiluminescence (Immobilon Western HRP Substrate, WBKLS0500, Millipore).

### i-POND assay

After addition of tetracycline (TET) to induce FLAG-Set8^PIPmut^ expression and eventually G1/S synchronization with thymidine before released into S-phase, 10^8^ cells were incubated with 10 µM EdU for 20 minutes to label ongoing replication forks before fixation with 2% formaldehyde for 20 minutes at a concentration of 2×10^6^ cells/ml. Formaldehyde quenching was done by adding 125 mM Glycine for 5 minutes. Cells were then permeabilized for 10 minutes on ice with PBS containing 0.25% Triton. Samples were then incubated in PBS solution containing 2 mM CUSO4 and 2 mM BTTP for 60 min at the concentration of 20×10^6^ cells/ml, before EdU conjugation to Biotin (Click reaction) by adding 10 mM Sodium-L-Ascorbate, 20 µM Biotin-TEG (Sigma-Aldrich) for 30 minutes on a rotating wheel (Click reaction). For “No Click” condition, Biotin-TEG was replaced by DMSO. Cells were then lysed in LB3JD Lysis Buffer (10 mM Hepes, pH 8, 100 mM NaCl, 500 mM EDTA pH 8, 100 mM EGTA pH 8, 0.2% SDS, 0.1% N-Lauroylsarcosine supplemented with 1mM PMSF and protease inhibitors (Roche)) at the concentration of 4×10^7^ cells/ml. Fixed chromatin was sheared using Bioruptor Pico (Diagenode) for 15 min or using the EpiShearTM Probe Sonicator probe. Samples were clarified by centrifugation at 15,000 rpm for 10 minutes. Aliquots corresponding to 1/100 of the final volume were saved for chromatin fragmentation analysis. 50 µl of streptavidin magnetic beads (Ademtech) were added to the EdU/Biotin-TEG-labeled chromatin and samples were incubated for 16-20h on a rotating wheel at 4°C in the dark. Beads were washed with LB3JD Lysis Buffer, then with 500 mM NaCl and twice again with LB3JD Lysis Buffer. Finally, before immunoblot analysis, purified DNA/protein complexes were eluted from beads and denatured with 50-70 µl of denaturation buffer (125 mM Tris-HCl pH6.8, 5% Glycerol, 2% SDS, 100 mM DTT, 2% β-mercaptoethanol) by heating at 95° C for 30 minutes with vigorous shaking. In parallel, one aliquot of total extract was denatured under the same conditions after adding of denaturation buffer at a 1:1 ratio. To verify chromatin fragmentation, aliquot was two times diluted in SDS buffer (50 mM Tris-HCl pH8, 1 mM EDTA pH8, 2% SDS) then heated to 65°C overnight with vigorous shacking. Aliquot was diluted in TE Buffer (50 mM Tris-HCl pH8, 100 mM NaCl, 1 mM EDTA) and treated with RNAse A (200 µg/ml, 45 min at 37°C) and with proteinase K (20 μg/ml, 30 min. at 37°C and 30 min. at 55°C). DNA was then purified with phenol/chloroform (Sigma-Aldrich) followed by ethanol/sodium acetate precipitation. DNA was analyzed on 2% agarose gel.

## LEGENDS OF SUPPLEMENTARY FIGURES

**Figure S1: FLAG-SET8^PIPmut^ expression during S-phase inhibits homologous recombination**. (**A**) bar-plot representing the percentages of EdU positive nuclei that showed BARD1, BRCA1, RAD51 and/or 53BP1 foci as indicated in control (−Tet) and tetracycline (+ Tet) treated FLAG-Set8^PIPmut^ cells. Data = mean ± s.d., n=3. ** p<0.01; *** p<0.001 (t-test). number of nuclei >100 per experiment. (**B**). Scattered box-plot representing the 53BP1 foci area in EdU positive nuclei from control (−Tet) and tetracyclin-treated (+ Tet) FLAG-SET8^PIPmut^ cells as in A. ** p<0.01; *** p<0.001. Inside the box-plot graphs, the thick line represents the median, the limit of the boxes corresponds to the 0.25 and 0.75 quartiles with whiskers showing values 1.5 times the interquartile range.n ≥ 3. *** p<0.001 (t-test), number of nuclei >100 per experiment. (**C**) Left panel is a scattered box-plot representing the number of γH2AX foci per EdU positive nucleus from control (−Tet) and FLAG-Set8^PIPmut^ expressing cells (+ Tet). Right panel is a box-plot representing the γH2AX foci area in EdU positive nuclei from control (−Tet) and FLAG-SET8^PIPmut^ expressing cells (+ Tet). *** p<0.001. Inside the box-plot graphs, the thick line represents the median, the limit of the boxes corresponds to the 0.25 and 0.75 quartiles with whiskers showing values 1.5 times the interquartile range. n≥3. *** p<0.001 (t-test). number of nuclei >100 per experiment.

**Figure S2: 53BP1 depletion did not rescue the loss of replication-coupled focal accumulation of BARD1 upon SET8 stabilization.** (**A**) Left panels are representative images showing staining of BARD1 and 53BP1 in EdU-positive nuclei from control and 53BP1 siRNA treated cells that were subsequent induced (+ Tet) or not (−Tet) for the expression of the FLAG-Set8^PIPmut^. Scale bar = 10 µm. Right panel is a representative scattered box-plot of the number of BARD1 foci per EdU positive nucleus from the same cells as above. *** p<0.001. number of analyzed nuclei = 100 per experiment (n=3). (**B**) Representative immunoblots showing the localization of BARD1 and 53BP1 in soluble cytoplasmic (S1) and nuclei (S2) fractions and in chromatin-enriched (P3) fraction in control and 53BP1 SiRNA-treated cells that were released into S phase after G1/S synchronization with thymidine treatment. Immunoblots with anti-histone 3 serves as control of the chromatin-enriched (P3) fraction.

**Figure S3: Levels of H4K20me states in FLAG-SET8^PIPmut^ U2OS cells treated with the SUV4-20H inhibitor A196 for 24 hours.** Immunoblots showing mono-, di- and trimethylation states levels of H4K20, and FLAG-SET8^PIPmut^ expression in FLAG-SET8^PIPmut^ U2OS cells co-treated or not with tetracyline and/or the SUV4-20H inhibitor A196 for 24 hours. Histone H4 and Tubulin were used as loading controls.

**Figure S4: Prolonged inhibition of SUV4-20H enzymes did not prevent 53BP1 focal accumulation and did not restore HR foci in FLAG-SET8^PIPmut^ cells**. (**A**). immunblots showing mono-, di- and trimethylation states levels of H4K20, and FLAG-SET8^PIPmut^ expression in control (−Tet) and FLAG-SET8^PIPmut^ (+Tet) cells that were treated (A196) or not (DMSO) with 10 µM of the SUV4-20H inhibitor A196 during 6 days. Histone 4 and Tubulin were used as loading controls. (**B**) Representative images showing staining of RAD51 and 53BP1 in EdU positive nuclei from control (−Tet) and FLAG-SET8^PIPmut^ expressing cells (+ Tet), and which were previously treated or not with 10mM of the SUV4-20H inhibitor A196 during 6 days. Scale bar = 10 µm. (**C**) Scattered box-plots representing the number of BARD1, BRCA1, 53BP1 or RAD51 foci per EdU positive nucleus from control (−Tet) and FLAG-SET8^PIPmut^ replicating cells (+ Tet) that were previously treated (A196) or not (DMSO) with 10mM of A196 during 6 days. Inside the box-plot graphs, the thick line represents the median, the limit of the boxes corresponds to the 0.25 and 0.75 quartiles with whiskers showing values 1.5 times the interquartile range. n=3. * p<0.05; **p<0.001 (t-test). number of nuclei >100 per experiment.

**Figure S5: RNF168 recruitment on replicated chromatin upon SET8 stabilization.** (**A**) Bar-plot representing the percentages of EdU positive nuclei displaying RNF168 and/or 53BP1 nuclear foci in control (−Tet) and tetracycline FLAG-SET8^PIPmut^ (+Tet) cells. Data = mean ± s.d., n =3. *p<0.05; ***p<0.001 (t-test). number of nuclei >100 per experiment **(B)** Scattered box-plot representing the number of RNF168 foci per EdU positive nucleus from control and 53BP1 depleted tetracylcine-inducible FLAG-SET8^PIPmut^ cells that were subsequent induced (+ Tet) or not (−Tet) for the expression of the FLAG-SET8^PIPmut^ protein for 24 hours. the thick line represents the median, the limit of the boxes corresponds to the 0.25 and 0.75 quartiles with whiskers showing values 1.5 times the interquartile range. ***p<0.001 **(C)** Scattered box-plots representing the number or RNF168 foci per EdU positive nucleus of Tet-inducible FLAG-SET8^PIPmut^ cells untreated (−Tet) or treated with tetracycline (+Tet) in presence or not of the Suv4-20h pharmacological inhibitor A196 (10 µM, 24h). The thick line represents the median, the limit of the boxes corresponds to the 0.25 and 0.75 quartiles with whiskers showing values 1.5 times the interquartile range. * p<0.05; ***p<0.001. Number of nuclei per condition and per experiment > 100. **(D)** Immunoblots showing the levels of RNF168 and FLAG-SET8^PIPmut^ protein in control (−Tet) and SET8^PIPmut^ (+ Tet) cells that were previously transfected with control or RNF168 siRNA two days before tetracycline treatment. Tubulin was used as loading control.

**Figure S6: Characterization of the camptothecin-induced DNA damage responses in control and FLAG-SET8^PIPmut^ expressing cells**. (**A)**. Western-blot showing the expression and phosphorylation levels of indicated proteins in control (−Tet) and FLAG-SET8^PIPmut^ (+ Tet) cells, untreated (DMSO) or treated with 5 nM or 25 nM of camptothecin during 2 hours. Tubulin was used as loading control. (**B)** Left panels are representative images of RNF168 and 53BP1 staining in EdU-positive nuclei from untreated (−Tet) and FLAG-Set8^PIPmut^ (+Tet) cells, two hours after exposure of not with camptothecin as indicated. Right panel is a scattered box-plot representing the number of RNF168 foci per EdU-positive nucleus from untreated (−Tet) and FLAG-Set8^PIPmut^ (+Tet) cells after exposure of not with camptothecin as indicated. Inside the box-plot graphs, the thick line represents the median, the limit of the boxes corresponds to the 0.25 and 0.75 quartiles with whiskers showing values 1.5 times the interquartile range. ***p<0.001. Number of nuclei per condition and per experiment > 100. n=3 (**C**). Left panels are representative images of the RPA70 and 53BP1 staining in control (−Tet) or FLAG-SET8^PIPmut^ (+ Tet) cells and that were untreated (DMSO) or treated with 25 nM of camptothecin during 2 hours. Scale bar = 10 µm. Right panel is a scattered box-plot representing the number of RPA foci per EdU positive nucleus from control (−Tet) and FLAG-SET8^PIPmut^ expressing cells (+ Tet) treated with 25 nM of camptothecin during 2 hours. n=3. number of nuclei >100 per experiment. (**D**) representative scattered box-plots of the number of RAD51 and BARD1 foci per EdU positive nucleus from control and 53BP1 siRNA treated cells that were subsequent induced (+ Tet) or not (−Tet) for the expression of the FLAG-Set8^PIPmut^ and treated with CPT (5 nM); number of analyzed nuclei = 50 per experiment (n=3).

**Figure S7: G1/S synchronization followed by SET8 depletion prevents the switch from H4K20me0 to H4K20me1 after DNA replication.** Upper panel is a cartoon showing the synchronization protocol. U2OS were synchronized with double thymidine block at G1/S transition, control and SET8 siRNA transfected 12h before G1/S release. Cells were then collected at different time points and cell-cycle progression was analyzed by FACS (lower panels). Proteins levels at each time points were analyzed by immunoblots (right panels).

## ACKNOWLEDGMENTS

This work was supported by Agence Nationale pour la Recherche (ANR-20-CE12-000 - LysMeth to E.J.), SIRIC Montpellier Cancer grant (to E.J.) and Ligue Contre Le Cancer Languedoc Roussillon (520027FF to CG). Institutional Support was provided by the Institut National de la Santé et de la Recherche Médicale (INSERM), by the Centre National de la Recherche Scientifique (CNRS). Y.P. was supported by a fellowship from Ligue Nationale Contre le Cancer (LNCC) FRM (Fondation d ela Recherche Médicale), F.A. by a fellowship from Université Montpelllier (UM)/CNRS-Liban and by LNCC. We thank Florence Cammas for helpful conceptual advices.

## AUTHORS CONTRIBUTION

YP, CG and EJ designed the project, CG and EJ supervised the research. YP, CG, JP, ML, VB, FA, DL and EJ performed the experiments. YP, CG, FA, VB and EJ analyzed the data. CR, VB and DL provided conceptual advices. ML, VB, YP, CG and EJ wrote the paper with the help of VB, YP and CG. EJ, FA, YP, CG, VB and CR contributed to revision and manuscript editing.

## REFERENCES

[1] Xu Y, Xu D. Repair pathway choice for double-strand breaks. Wu Q, editor. Essays Biochem. 2020;64(5):765–777. doi: 10.1042/EBC20200007

[2] Scully R, Panday A, Elango R, et al. DNA double-strand break repair-pathway choice in somatic mammalian cells. Nat Rev Mol Cell Biol. 2019;20(11):698–714. doi: 10.1038/s41580-019-0152-0.

[3] Hustedt N, Durocher D. The control of DNA repair by the cell cycle. Nat Cell Biol 2016, 19(1):1–9. doi: 10.1038/ncb3452.

[4] Kolinjivadi AM, Sannino V, de Antoni A, et al. Moonlighting at replication forks - a new life for homologous recombination proteins BRCA1, BRCA2 and RAD51. FEBS Lett. 2017; 591(8):1083–1100. doi: 10.1002/1873-3468.12556.

[5] Toh M, Ngeow J. Homologous Recombination Deficiency: Cancer Predispositions and Treatment Implications. The Oncologist 2021;26(9):e1526–e1537. doi: 10.1002/onco.13829

[6] Jeggo PA, Löbrich M. How cancer cells hijack DNA double-strand break repair pathways to gain genomic instability. Biochem J 2015;471(1):1–11. doi: 10.1042/BJ20150582

[7] Fradet-Turcotte A, Canny MD, Escribano-Díaz C, et al. 53BP1 is a reader of the DNA-damage-induced H2A Lys 15 ubiquitin mark. Nature. 2013;499(7456):50–54. doi: 10.1038/nature12318

[8] Tarsounas M, Sung P. The antitumorigenic roles of BRCA1–BARD1 in DNA repair and replication. Nat Rev Mol Cell Biol 2020;21(5):284–299. doi: 10.1038/s41580-020-0218-z

[9] Michelena J, Pellegrino S, Spegg V, et al. Replicated chromatin curtails 53BP1 recruitment in BRCA1-proficient and BRCA1-deficient cells. Life Sci Alliance. 2021;4(6):e202101023. doi: 10.26508/lsa.202101023

[10] Botuyan MV, Lee J, Ward IM, et al. Structural basis for the methylation state-specific recognition of histone H4-K20 by 53BP1 and Crb2 in DNA repair. Cell. 2006;127(7):1361–1373. doi: 10.1016/j.cell.2006.10.043.

[11] Svobodová Kovaříková A, Legartová S, Krejčí J, et al. H3K9me3 and H4K20me3 represent the epigenetic landscape for 53BP1 binding to DNA lesions. Aging. 2018;10(10):2585–2605. doi: 10.18632/aging.101572

[12] Nakamura K, Saredi G, Becker JR, et al. H4K20me0 recognition by BRCA1–BARD1 directs homologous recombination to sister chromatids. Nat Cell Biol. 2019;21(3):311–318. doi: 10.1038/s41556-019-0282-9.

[13] Witus SR, Zhao W, Brzovic PS, et al. BRCA1/BARD1 is a nucleosome reader and writer. Trends Biochem Sci. 2022;47(7):582–595. doi: 10.1016/j.tibs.2022.03.001.

[14] Alabert C, Barth TK, Reverón-Gómez N, et al. Two distinct modes for propagation of histone PTMs across the cell cycle. Genes Dev. 2015; 29 (6):585–590. doi: 10.1101/gad.256354.114.

[15] Cao X, Chen Y, Wu B, et al. Histone H4K20 Demethylation by Two hHR23 Proteins. Cell Rep. 2020;30(12):4152–4164.e6. doi: 10.1016/j.celrep.2020.03.001.

[16] Liu W, Tanasa B, Tyurina OV, et al. PHF8 mediates histone H4 lysine 20 demethylation events involved in cell cycle progression. Nature. 2010; 466(7305):508–512. doi: 10.1038/nature09272.

[17] Saredi G, Huang H, Hammond CM, et al. H4K20me0 marks post-replicative chromatin and recruits the TONSL–MMS22L DNA repair complex. Nature. 2016;534(7609):714–718. doi: 10.1038/nature18312.

[18] Beck DB, Burton A, Oda H, et al. The role of PR-Set7 in replication licensing depends on Suv4-20h. Genes Dev. 2012;26(23):2580–2589. doi: 10.1101/gad.195636.112

[19] Brustel J, Tardat M, Kirsh O, et al. Coupling mitosis to DNA replication: the emerging role of the histone H4-lysine 20 methyltransferase PR-Set7. Trends Cell Biol. 2011;21(8):452–460. doi: 10.1016/j.tcb.2011.04.006.

[20] Jørgensen S, Schotta G, Sørensen CS. Histone H4 lysine 20 methylation: key player in epigenetic regulation of genomic integrity. Nucleic Acids Res. 2013;41(5):2797–2806. doi: 10.1093/nar/gkt012

[21] Centore RC, Havens CG, Manning AL, et al. CRL4(Cdt2)-mediated destruction of the histone methyltransferase Set8 prevents premature chromatin compaction in S phase. Mol Cell. 2010;40(1):22–33. doi: 10.1016/j.molcel.2010.09.015.

[22] Oda H, Hübner MR, Beck DB, et al. Regulation of the histone H4 monomethylase PR-Set7 by CRL4(Cdt2)-mediated PCNA-dependent degradation during DNA damage. Mol Cell. 2010;40(3):364–376. doi: 10.1016/j.molcel.2010.10.011.

[23] Tardat M, Brustel J, Kirsh O, et al. The histone H4 Lys 20 methyltransferase PR-Set7 regulates replication origins in mammalian cells. Nat Cell Biol. 2010;12(11):1086–1093. doi: 10.1038/ncb2113

[24] Rice JC, Nishioka K, Sarma K, et al. Mitotic-specific methylation of histone H4 Lys 20 follows increased PR-Set7 expression and its localization to mitotic chromosomes. Genes Dev. 2002;16(17):2225–2230. doi: 10.1101/gad.1014902.

[25] Dungrawala H, Cortez D. Purification of proteins on newly synthesized DNA using iPOND. Methods Mol Biol Clifton NJ. 2015;1228:123–131. doi: 10.1007/978-1-4939-1680-1_10.

[26] Aze A, Zhou JC, Costa A, et al. DNA replication and homologous recombination factors: acting together to maintain genome stability. Chromosoma. 2013;122(5):401–413. doi: 10.1007/s00412-013-0411-3.

[27] Chapman JR, Sossick AJ, Boulton SJ, et al. BRCA1-associated exclusion of 53BP1 from DNA damage sites underlies temporal control of DNA repair. J Cell Sci. 2012;125(15):3529–3534. doi: 10.1242/jcs.105353

[28] Bhowmick R, Lerdrup M, Gadi SA, et al. RAD51 protects human cells from transcription-replication conflicts. Mol Cell. 2022;82(18):3366–3381.e9. doi: 10.1016/j.molcel.2022.07.010

[29] Costanzo V. Brca2, Rad51 and Mre11: Performing balancing acts on replication forks. DNA Repair. 2011;10(10):1060–1065. doi: 10.1016/j.dnarep.2011.07.009.

[30] Feu S, Unzueta F, Ercilla A, et al. RAD51 is a druggable target that sustains replication fork progression upon DNA replication stress. Cotterill S, editor. PLOS ONE. 2022;17(8):e0266645. doi: 10.1371/journal.pone.0266645

[31] Pellegrino S, Michelena J, Teloni F, et al. Replication-Coupled Dilution of H4K20me2 Guides 53BP1 to Pre-replicative Chromatin. Cell Rep. 2017;19(9):1819–1831. doi: 10.1016/j.celrep.2017.05.016.

[32] Krais JJ, Wang Y, Patel P, et al. RNF168-mediated localization of BARD1 recruits the BRCA1-PALB2 complex to DNA damage. Nat Commun. 2021;12(1):5016. doi: 10.1038/s41467-021-25346-4.

[33] Zong D, Adam S, Wang Y, et al. BRCA1 Haploinsufficiency Is Masked by RNF168-Mediated Chromatin Ubiquitylation. Mol Cell. 2019;73(6):1267–1281.e7. doi: 10.1016/j.molcel.2018.12.010.

[34] Zong D, Callén E, Pegoraro G, et al. Ectopic expression of RNF168 and 53BP1 increases mutagenic but not physiological non-homologous end joining. Nucleic Acids Res. 2015;43(10):4950–4961. doi: 10.1093/nar/gkv336.

[35] Dulev S, Lin S, Liu Q, et al. SET8 localization to chromatin flanking DNA damage is dependent on RNF168 ubiquitin ligase. Cell Cycle. 2020;19(1):15–23. doi: 10.1080/15384101.2019.1690231.

[36] Lu X, Xu M, Zhu Q, et al. RNF8-ubiquitinated KMT5A is required for RNF168-induced H2A ubiquitination in response to DNA damage. FASEB J. 2021;35(4). doi: 10.1096/fj.202002234R

[37] Becker JR, Clifford G, Bonnet C, et al. BARD1 reads H2A lysine 15 ubiquitination to direct homologous recombination. Nature. 2021;596(7872):433–437. doi: 10.1038/s41586-021-03776-w

[38] Han S, Lim KS, Blackburn BJ, et al. The Potential of Topoisomerase Inhibitor-Based Antibody–Drug Conjugates. Pharmaceutics 2022;14(8):1707. doi: 10.3390/pharmaceutics14081707

[39] Shoaib M, Walter D, Gillespie PJ, et al. Histone H4K20 methylation mediated chromatin compaction threshold ensures genome integrity by limiting DNA replication licensing. Nat Commun. 2018;9(1):3704. doi: 10.1038/s41467-018-06066-8

[40] Callen E, Zong D, Wu W, et al. 53BP1 Enforces Distinct Pre- and Post-resection Blocks on Homologous Recombination. Mol Cell. 2020;77(1):26–38.e7. doi: 10.1016/j.molcel.2019.09.024.

[41] Shi X, Kachirskaia I, Yamaguchi H, et al. Modulation of p53 function by SET8-mediated methylation at lysine 382. Mol Cell. 2007;27(4):636–646. doi: 10.1016/j.molcel.2007.07.012.

[42] Takawa M, Cho H-S, Hayami S, et al. Histone lysine methyltransferase SETD8 promotes carcinogenesis by deregulating PCNA expression. Cancer Res. 2012;72(13):3217–3227. doi: 10.1158/0008-5472.CAN-11-3701.

[43] Bromberg KD, Mitchell TRH, Upadhyay AK, et al. The SUV4-20 inhibitor A-196 verifies a role for epigenetics in genomic integrity. Nat Chem Biol. 2017;13(3):317–324. doi: 10.1038/nchembio.2282

[44] Schotta G, Sengupta R, Kubicek S, et al. A chromatin-wide transition to H4K20 monomethylation impairs genome integrity and programmed DNA rearrangements in the mouse. Genes Dev. 2008;22(15):2048–2061. doi: 10.1101/gad.476008.

[45] Bunting SF, Callén E, Wong N, et al. 53BP1 Inhibits Homologous Recombination in Brca1-Deficient Cells by Blocking Resection of DNA Breaks. Cell. 2010;141(2):243–254. doi: 10.1016/j.cell.2010.03.012

[46] Irminger-Finger I, Leung WC. BRCA1-dependent and independent functions of BARD1. Int J Biochem Cell Biol. 2002;34(6):582–587. doi: 10.1016/s1357-2725(01)00161-3.

[47] Savage KI, Harkin DP. BRCA1, a “complex” protein involved in the maintenance of genomic stability. FEBS J. 2015;282(4):630–646. doi: 10.1111/febs.13150.

[48] Jørgensen S, Elvers I, Trelle MB, et al. The histone methyltransferase SET8 is required for S-phase progression. J Cell Biol. 2007;179(7):1337–1345. doi: 10.1083/jcb.200706150.

[49] Escribano-Díaz C, Orthwein A, Fradet-Turcotte A, et al. A cell cycle-dependent regulatory circuit composed of 53BP1-RIF1 and BRCA1-CtIP controls DNA repair pathway choice. Mol Cell. 2013;49(5):872–883. doi: 10.1016/j.molcel.2013.01.001

[50] Orthwein A, Noordermeer SM, Wilson MD, et al. A mechanism for the suppression of homologous recombination in G1 cells. Nature. 2015;528(7582):422–426. doi: 10.1038/nature16142.

[51] Yilmaz D, Furst A, Meaburn K, et al. Activation of homologous recombination in G1 preserves centromeric integrity. Nature. 2021;600(7890):748–753. doi: 10.1038/s41586-021-04200-z

[52] Liao T, Wang Y-J, Hu J-Q, et al. Histone methyltransferase KMT5A gene modulates oncogenesis and lipid metabolism of papillary thyroid cancer in vitro. Oncol Rep. 2018;39(5):2185–2192. doi: 10.3892/or.2018.6295

[53] Milite C, Feoli A, Viviano M, et al. The emerging role of lysine methyltransferase SETD8 in human diseases. Clin Epigenetics. 2016;8:102. doi: 10.1186/s13148-016-0268-4.

[54] Yang C, Wang K, Zhou Y, et al. Histone lysine methyltransferase SET8 is a novel therapeutic target for cancer treatment. Drug Discov Today. 2021;26(10):2423–2430. doi: 10.1016/j.drudis.2021.05.004

[55] Hernández-Reyes Y, Paz-Cabrera MC, Freire R, et al. USP29 Deubiquitinates SETD8 and Regulates DNA Damage-Induced H4K20 Monomethylation and 53BP1 Focus Formation. Cells. 2022;11(16):2492. doi: 10.3390/cells11162492

[56] Veo B, Danis E, Pierce A, et al. Combined functional genomic and chemical screens identify SETD8 as a therapeutic target in MYC-driven medulloblastoma. JCI Insight. 2019;4(1). doi: 10.1172/jci.insight.122933

[57] Veschi V, Liu Z, Voss TC, et al. Epigenetic siRNA and Chemical Screens Identify SETD8 Inhibition as a Therapeutic Strategy for p53 Activation in High-Risk Neuroblastoma. Cancer Cell. 2017;31(1):50–63. doi: 10.1016/j.ccell.2016.12.002.

[58] Tardat M, Murr R, Herceg Z, et al. PR-Set7-dependent lysine methylation ensures genome replication and stability through S phase. J Cell Biol. 2007;179(7):1413–1426. doi: 10.1083/jcb.200706179

