## Supplementary figures and images for "Cell-cycle dependent inhibition of BRCA1 signaling by the lysine methyltransferase SET8"

### supplementary figures

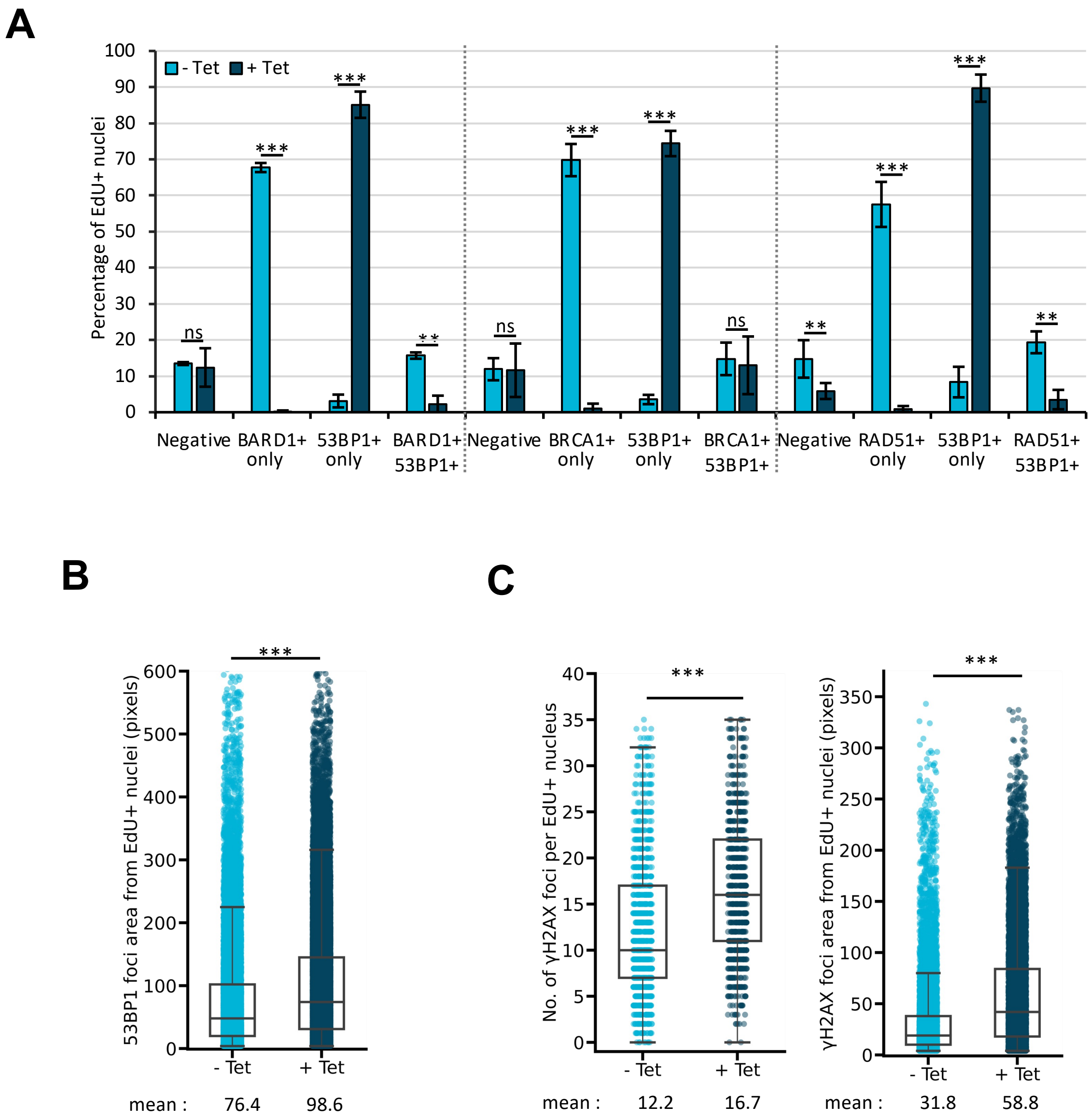

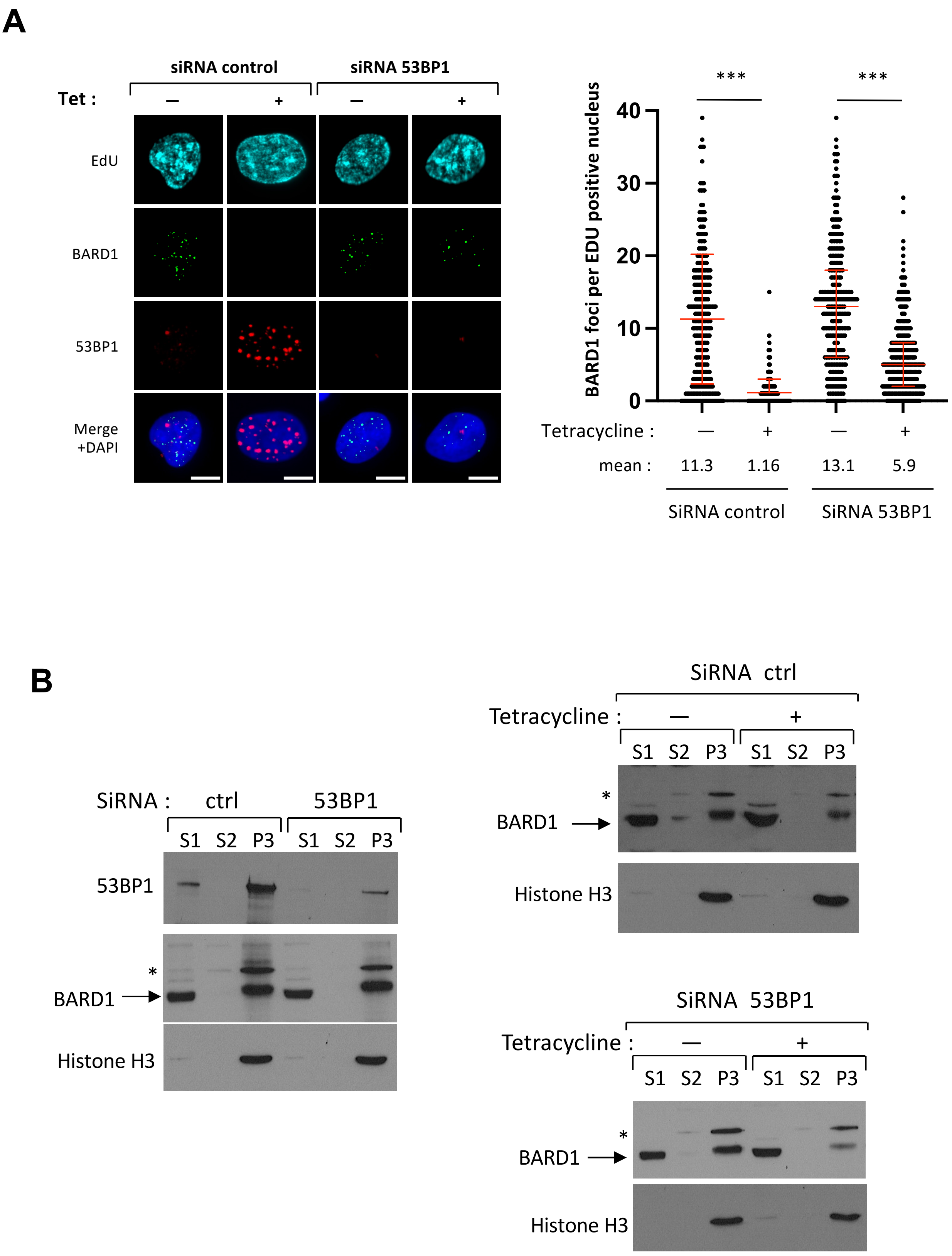

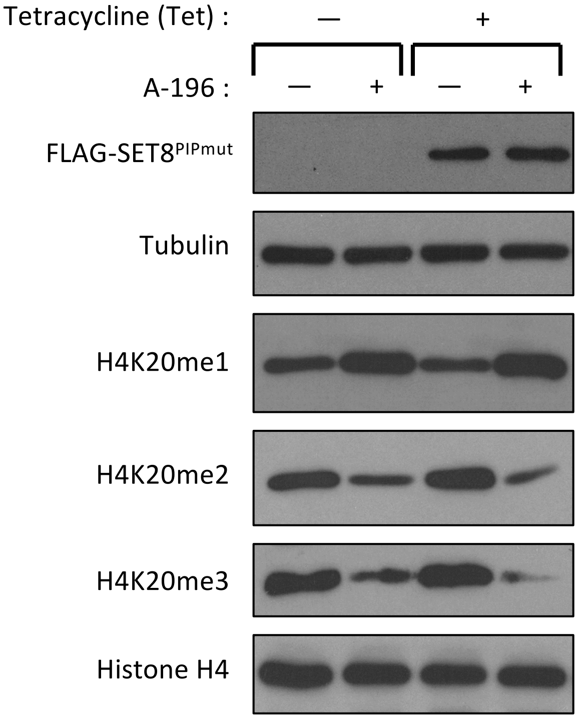

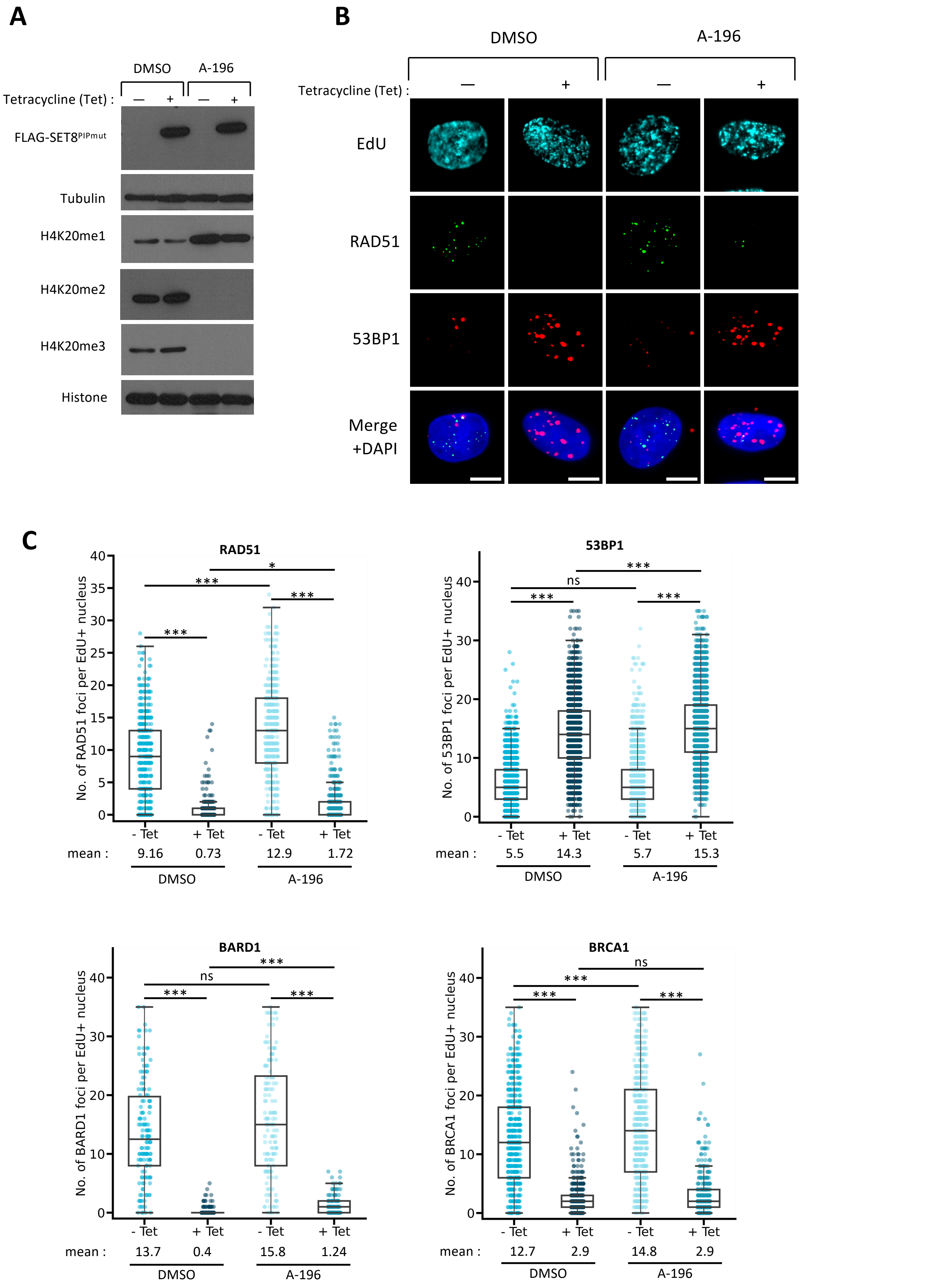

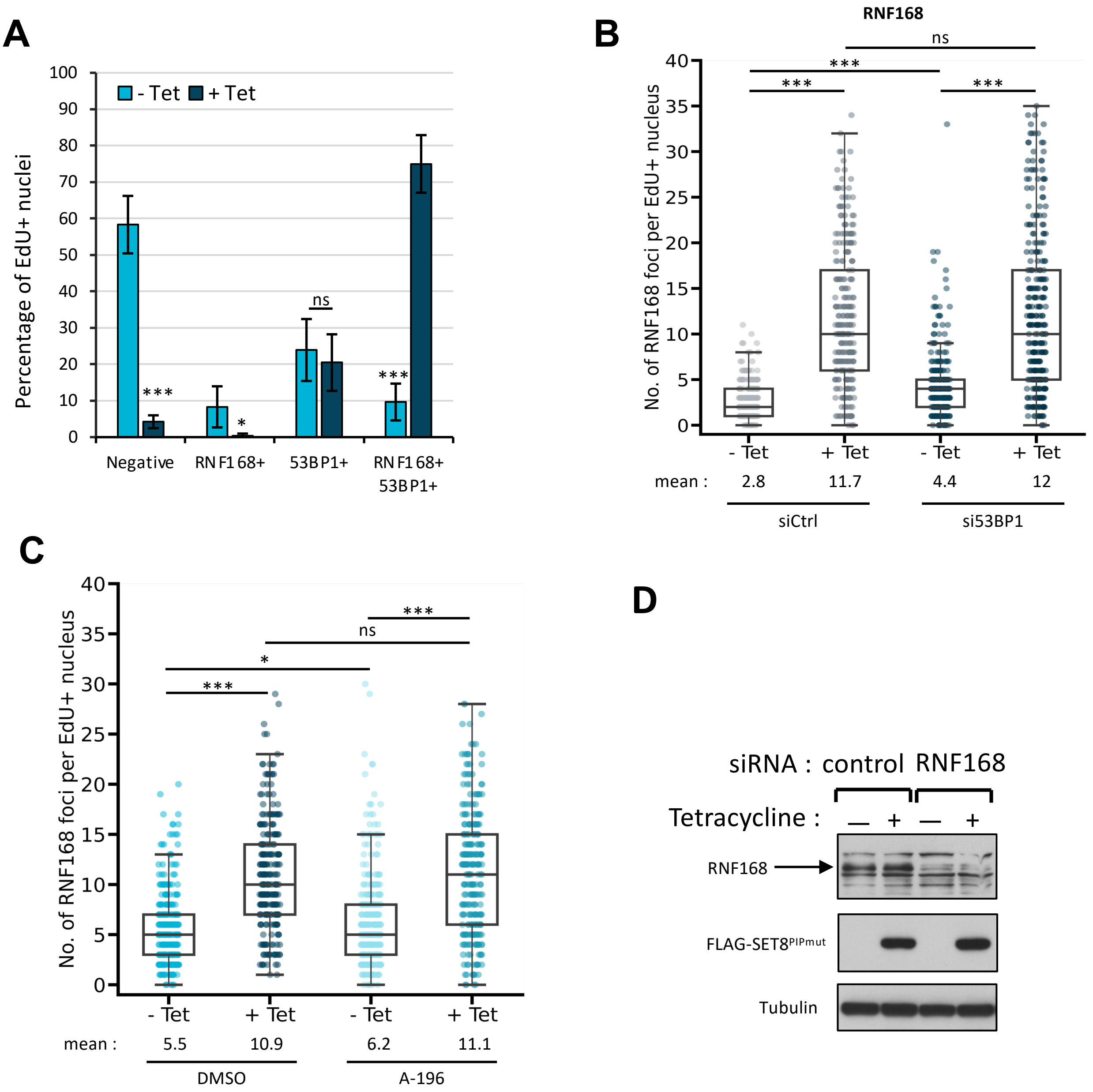

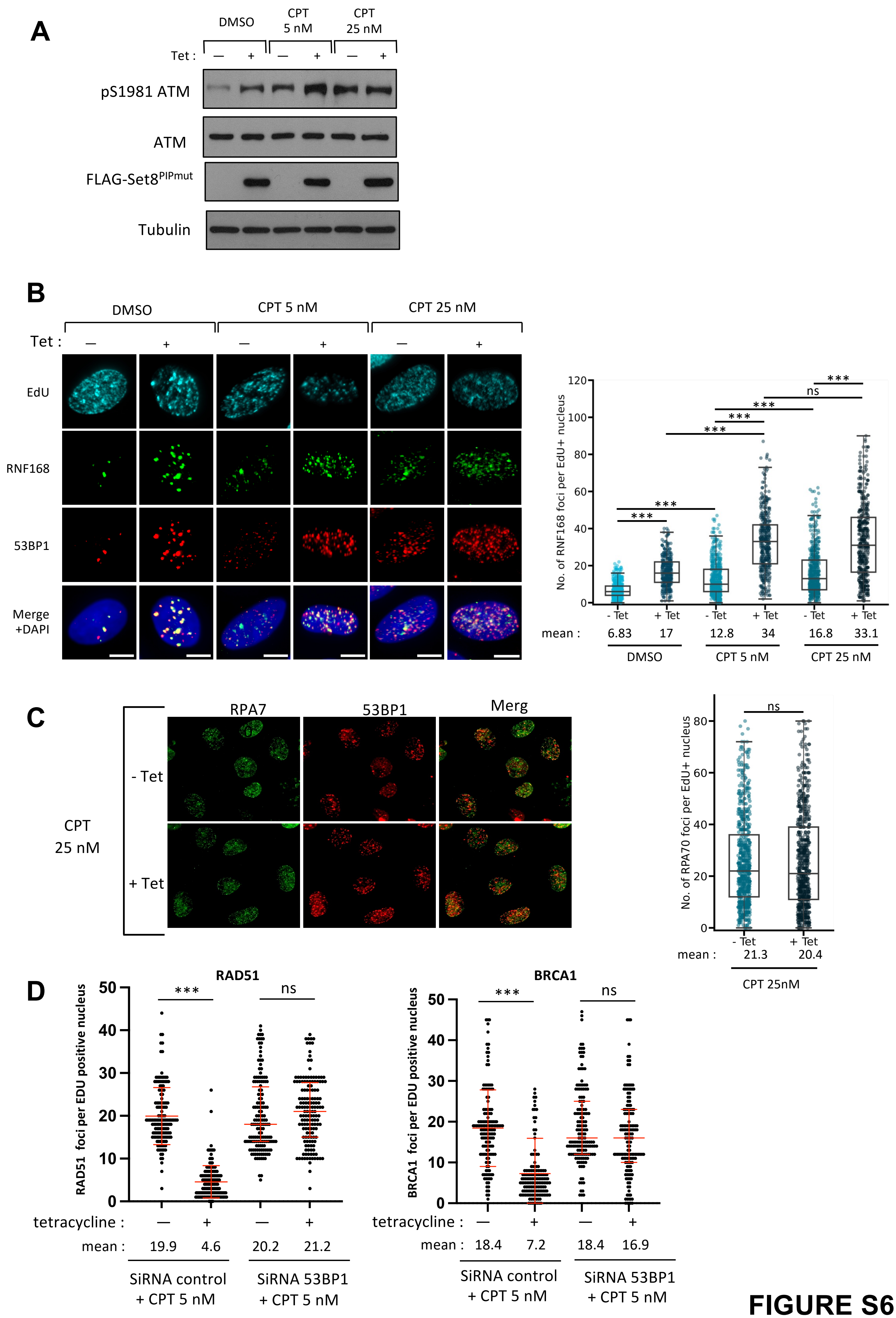

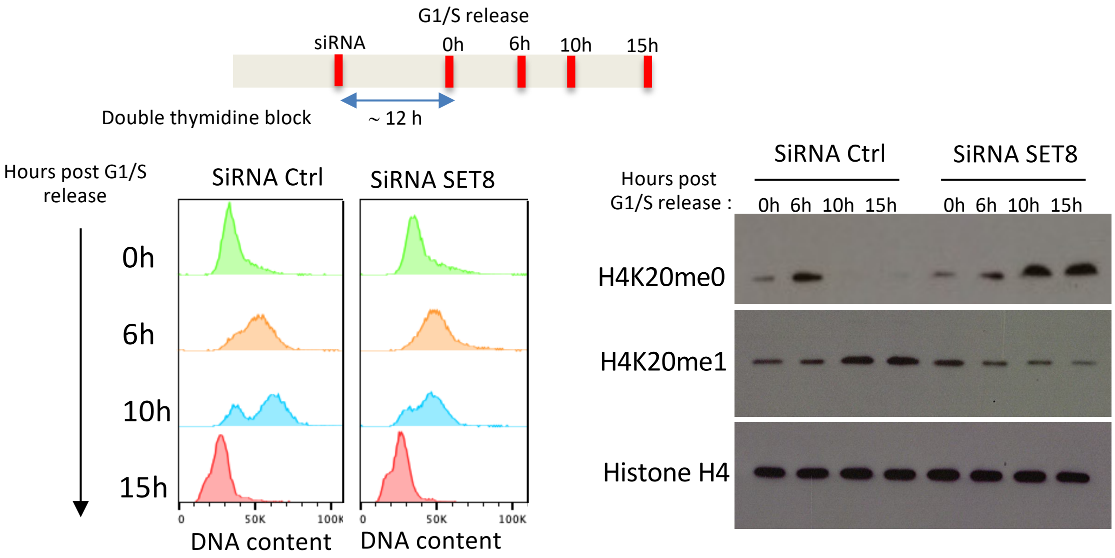
